# An Image Quality Transfer Technique for Localising Deep Brain Stimulation Targets

**DOI:** 10.1101/2024.03.18.584979

**Authors:** Ying-Qiu Zheng, Harith Akram, Zeju Li, Stephen M Smith, Saad Jbabdi

## Abstract

The ventral intermediate nucleus of the thalamus (Vim) is a well-established surgical target in functional neurosurgery for the treatment of tremor. As the structure lacks intrinsic contrast on conventional MRI sequences, targeting the Vim has predominantly relied on standardised Vim atlases which can fail to account for individual anatomical variability. To overcome this limitation, recent studies define the Vim using its structural connectivity profile generated via tractography. Although successful in accounting for individual variability, these connectivity-based methods are sensitive to variations in image acquisition and processing, and require high-quality diffusion imaging protocols which are usually not available in clinical settings. Here we propose a novel transfer learning approach to accurately target the Vim particularly on clinical-quality data. The approach transfers anatomical information from publicly-available high-quality datasets to a wide range of white matter connectivity features in low-quality data to augment inference on the Vim. We demonstrate that the approach can robustly and reliably identify Vim even with compromised data quality and is generalisable to datasets acquired with different protocols, outperforming previous surgical targeting methods. The approach is not limited to targeting Vim and can be adapted to other deep brain structures.

## I. INTRODUCTION

Functional neurosurgical techniques, including deep brain stimulation (DBS) and MR-guided focused ultrasound, have been successfully used for several decades to treat a range of neurological and psychiatric disorders. These techniques target specific neural circuits that are implicated in the pathophysiology of these disorders, allowing for modulation of circuit activity and resulting in the alleviation of symptoms that cannot be managed by medications alone. For instance, the ventral intermediate nucleus of the thalamus (Vim) is a well-established surgical target in DBS and stereotactic ablation for tremor in Parkinson’s Disease (PD), essential tremor, and multiple sclerosis [18, 23]. The Vim is a wedge-shaped nucleus located inferiorly within the ventrolateral nucleus [9, 36], and is one of the motor thalamic nuclei that play a critical role in tremor circuitry. Classical tract-tracing and immuno-histochemical studies have revealed that the Vim receives efferent fibres from the dentate nucleus of the contralateral cerebellum via the superior cerebellar peduncle at the midbrain level [29], and projects primarily to the ipsilateral primary motor cortex (M1) [43] with minor projections to the supplementary motor area and premotor cortex [53]. Collectively, the tract connecting the contralateral dentate nucleus to M1 via the Vim in the thalamus forms the dentato-thalamo-cortical pathway and plays a central role in tremor circuitry [10, 17, 21, 29, 34, 47].

Accurately targeting the Vim has remained a significant challenge due to the lack of intrinsic contrast in conventional MRI sequences to distinguish the nucleus from neighbouring structures and tissues. Traditional stereotactic targeting of the structure has relied on standardised atlases/coordinates adapted to individual subjects, aided by visible anatomical landmarks such as the anterior commissure and posterior commissure points [32]. Although stereotactic atlases provide a reproducible way to identify the Vim, they fail to account for inter-individual anatomical variations, which are not negligible in thalamic nuclei [13, 38, 42, 49]. To ensure efficacy of stereotactic surgery, patients are often required to stay awake in order to allow target confirmation, causing great patient discomfort and potential risks of intracranial haemorrhage leading to neurological deficit or even death [64].

To address this limitation, recent studies have attempted to more accurately identify the Vim in individual patients by leveraging its anatomical properties [1, 15, 24]. Using cutting-edge diffusion MRI (dMRI) techniques and tractography-based methods, these studies identify Vim by locating the thalamic region of maximum connectivity with both ipsilateral M1 and the contralateral dentate. Diffusion MRI enables the estimation of local fibre orientations, while tractography algorithms generate streamline samples that represent the underlying white matter pathway based on these local fibre orientations. This connectivity-driven approach can better explain the inter-individual anatomical variability of the nucleus, leading to improved efficacy of neurosurgical procedures [1]. However, reconstructing these connections usually requires state-of-the-art high angular resolution diffusion imaging, high spatial resolution with sufficient signal to noise, and advanced diffusion modelling techniques in order to resolve complex fibre configurations (e.g., crossing fibres), which are impractical in advanced-care clinical settings due to their prolonged acquisition time and higher acquisition/computational cost. As a result, lower-quality diffusion MRI techniques are still commonly used, which affects the reliability of the connectivity-driven approach as a proxy for surgical targeting. Furthermore, even with cutting-edge dMRI data and higher-order diffusion modelling, the Vim localised by the connectivity-driven approach has exhibited significant variations across different acquisition protocols (e.g., b-values, spatial and angular resolutions) and processing pipelines (e.g., diffusion signal modelling and tractography algorithm parameters) [15]. Therefore, these methods must be used with caution to ensure that they accurately reflect the true underlying anatomical variability rather than reflecting methodological confounds erroneously interpreted as variability [15].

Accurate localisation of the target nucleus is crucial for the efficacy of stereotactic operations. However, existing surgical targeting approaches have not yet been able to translate into a reliable clinical routine. To address this challenge, we propose a novel Image Quality Transfer (IQT) technique, HQ-augmentation, for reliably localising Vim, particularly on clinical-quality data. IQT is a computational technique in medical imaging that aims to enhance the quality of low-resolution or low-quality images by transferring information from a set of high-quality reference images [2, 3, 16, 58]. This is usually achieved by using machine learning algorithms to learn a mapping between low-quality images and their high-quality counterparts. Following this concept, the HQ-augmentation approach proposed here leverages anatomical information from publicly-available dMRI datasets, such as the Human Connectome Project (HCP), to guide Vim localisation in low-quality diffusion MRI datasets. It also exploits an augmented set of connectivity features with a wide range of brain regions to compensate for the compromised M1 and dentate white matter connectivity features in low-quality diffusion MRI (Figure 1). Specifically, we generate approximate locations of the Vim from high-quality diffusion MRI datasets and train the HQ-augmentation model to locate the nucleus in low-quality data that is most equivalent to the counterpart high-quality estimate, given its low-quality anatomical connectivity profiles with a wide range of brain regions. Our results demonstrate that the HQ-augmentation model outperforms existing alternatives, i.e., the atlas-defined approach and the connectivity-driven approach, on surrogate low-quality diffusion MRI data with much lower spatial and angular resolution, suggesting that the HQ-augmentation model is not only capable of accounting for the true anatomical variability but is also robust to the impact of data quality and methodological heterogeneity. Furthermore, the HQ-augmentation model can generalise to unseen low-quality diffusion MRI datasets collected with different acquisition protocols, such as the UK Biobank (UKB) dataset.

**FIG. 1.**
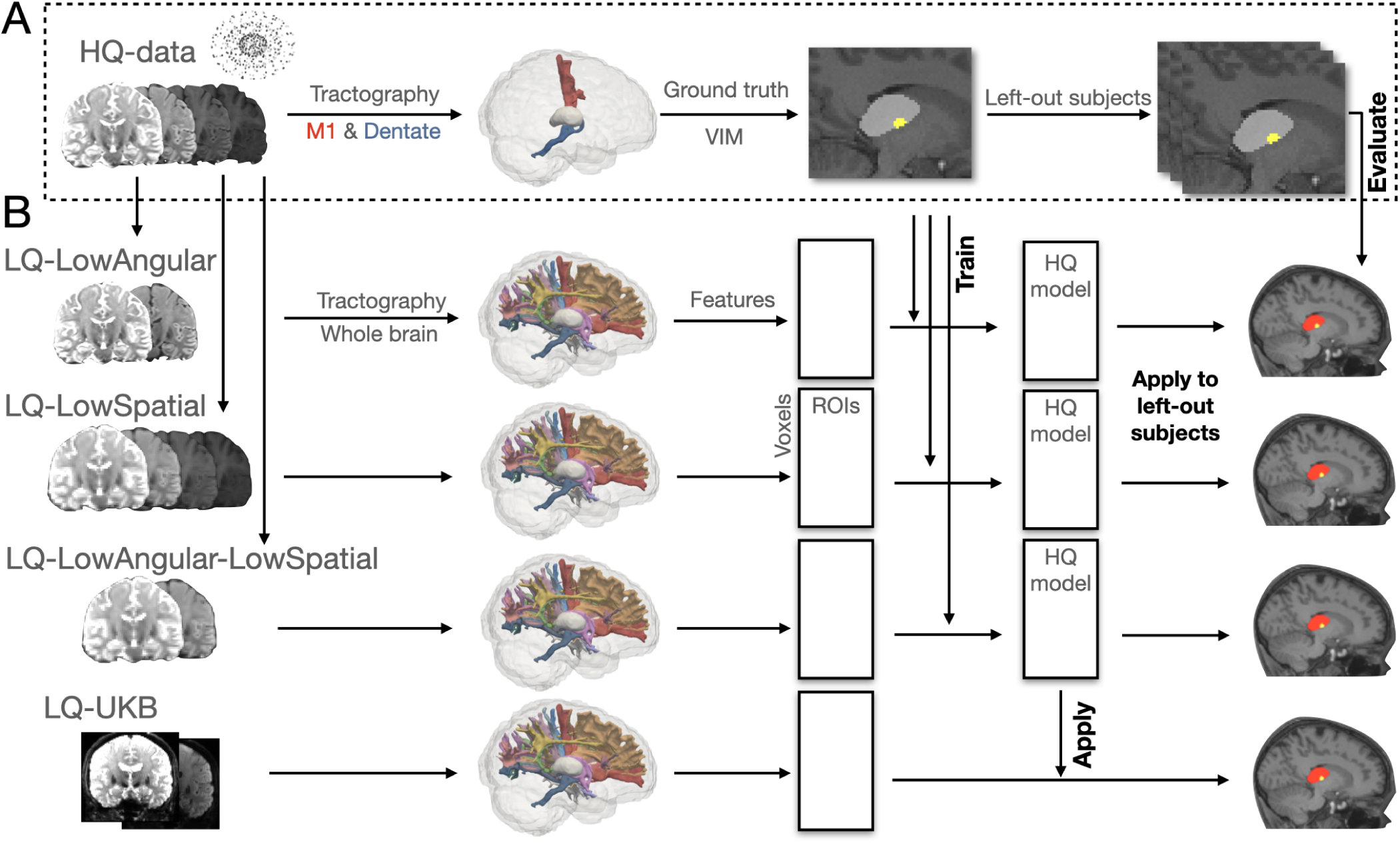
Illustration of the HQ-augmentation approach. **(A)** First, the HQ diffusion data were used to create “ground truth” Vim within the thalamic masks, based on its white matter connectivity with M1 and contralateral cerebellum. **(B)** Next, surrogate low-quality diffusion datasets were created by degrading the HQ datasets. The HQ-augmentation model was trained with the HCP HQ “ground truth” Vim as target labels and an extended set of HCP low-quality connectivity profiles as input features. The trained model was applied to unseen low-quality datasets (surrogate low-quality HCP and UK Biobank diffusion data) to generate Vim predictions, and evaluated against the corresponding HQ “ground truth”.

Finally, the HQ-augmentation model is not limited to targeting the Vim. Depending on specific symptoms, a range of brain areas have been revealed as effective surgical targets. Within the thalamus, for example, the anterior nucleus has been used as a DBS target to treat epilepsy [28, 37, 46, 51]; the medial dorsal nucleus is another DBS target to treat obssessive-compulsive disorder and major depressive disorder [33]. Outside of the thalamus, the subthalamic nucleus [14, 45, 52] and the globus pallidus internus [5, 50, 61] are common DBS targets for the treatment of Parkinson’s Disease; the hypothalamus has been explored for the treatment of cluster headaches, obesity, and other conditions. Our approach can be adapted to many other deep brain nuclei, using their literature-based white matter connectivity profiles. Overall, the HQ-augmentation serves as a better proxy for surgical targeting and has the potential to translate into a reliable clinical tool.

## II. MATERIALS AND METHODS

### A. Datasets and subjects

#### 1. HCP 3T minimally preprocessed MRI

We used 3T diffusion MRI data from the Human Connectome Project (HCP) [56, 59] as the high-quality dataset. The data were collected by the WU-Minn HCP consortium from a group of healthy young adults (n = 1,062, age range 22-36 years) who consented to participate under the approval of the Washington University in St Louis Institutional Review Board [59]. The minimally pre-processed T1-, T2-, and diffusion-weighted MRI scans were obtained from the HCP 2017 S1200 release (https://www.humanconnectome.org/study/hcp-young-adult/document/1200-subjects-data-release), which included 43 subjects who underwent repeated scanning. T1-and T2-weighted images were acquired with a customised Siemens 3T “Connectome Skyra” scanner at an isotropic spatial resolution of 0.7mm. Diffusion MRI data were collected with a monopolar diffusion-weighted (Stejskal-Tanner) spin-echo EPI sequence using the Siemens 3T “Connectome Skyra” scanner at an isotropic spatial resolution of 1.25mm. The acquisition included three shells (b-values = 1000, 2000, and 3000 s/mm^2^) and 90 unique diffusion directions per shell, acquired twice (total scan time 60 min per subject) [56]. As part of the HCP S1200 release, the data were minimally pre-processed [30] and aligned across modalities for each subject. Motion, susceptibility, and eddy current distortions in the diffusion MRI data were corrected [7, 8]. The data were linearly registered to the corresponding T1-weighted images using FSL’s FLIRT [40, 41]. Nonlinear transformations to the MNI152 standard space and their inverse warps were obtained using FSL’s FNIRT [6, 41]. The complete imaging protocols and pre-processing procedures can be found at https://www.humanconnectome.org/storage/app/media/documentation/s1200/HCP_S1200_Release_Reference_Manual.pdf.

#### 2. UK Biobank 3T minimally pre-processed MRI

We also used 3T MRI data from UK Biobank [48] in our analysis, as the overall quality of this data is more representative of what is typically acquired on clinical scanners. Our analysis included a total of 2,560 subjects who had been scanned twice (typically with a 2-year interval). T1-weighted images were acquired using a 3D MPRAGE acquisition at 1mm isotropic resolution, while T2-weighted images were acquired using a 3D SPACE sequence at a spatial resolution of 1.05×1×1mm. Diffusion MRI was performed at isotropic spatial resolution 2mm with two shells (b-values = 1000 and 2000 s/mm^2^), and 50 diffusion directions per shell (total scan time of approximately 6 minutes per subject). Additional information on the imaging protocols can be found in https://biobank.ctsu.ox.ac.uk/crystal/ crystal/docs/brain_mri.pdf. Pre-processing of the UK Biobank data included face removal, brain extraction, and registration across modalities and to MNI152 standard space [4]. The diffusion MRI images were also minimally pre-processed, with correction for motion, susceptibility and eddy current distortions [7, 8]. It is important to note that the (uncorrupted) UK Biobank 3T diffusion MRI may still be considered as a high-quality dataset. In the following analysis, we also derived a surrogate low-quality dataset from the UK Biobank dataset.

### B. Low-quality diffusion MRI datasets

To account for the varying data quality typically encountered in clinical contexts, we considered a range of low-quality datasets, including:

1. surrogate HCP low-quality diffusion MRI, obtained by degrading the original (minimally preprocessed) high-quality HCP 3T diffusion MRI. Three different forms of data corruption were explored: a dataset with reduced angular resolution, referred to as *LQ-LowAngular*; a dataset with decreased spatial resolution, denoted *LQ-LowSpatial*; and a dataset with reductions in both angular and spatial resolution, named *LQ-LowAngular-LowSpatial*.
2. surrogate UKB low-quality diffusion MRI, obtained by reducing the angular resolution of the original (minimally pre-processed) UKB 3T diffusion MRI, which we refer to as *LQ-UKB*;

The HQ-augmentation model was trained using paired high- and low-quality HCP data, specifically, the original HCP 3T diffusion MRI and its various lower-quality counterparts (*LQ-LowAngular*, *LQ-LowSpatial*, or *LQ-LowAngular-LowSpatial*). The paired high- and low-quality datasets provide examples for the model to learn the relationship between the low-quality image features and their corresponding high-quality counterparts, enabling the model to extract and transfer high-quality anatomical information to enhance low-quality data. As a result, the trained model is tailored to the specific type of low-quality dataset on which it is trained.

Validation and evaluation were carried out on both HCP and UKB datasets. The Vim derived from low-quality data was assessed against two versions of “ground truth”: the connectivity-driven Vim obtained from the high-quality data (referred to as HQ-Vim in the following text), and the atlas-defined Vim adapted to individual native space from the group-average Vim location.

#### 1. LQ-LowAngular

The *LQ-LowAngular* dataset was designed to resemble properties of single-shell diffusion MRI collected with fewer diffusion directions. To create this dataset, we discarded all volumes corresponding to bvals = 2000 and 3000 s/mm^2^ and only considered the single-shell at bvals = 1000 s/mm^2^ along with the b0 volumes. We then sampled 32 directions uniformly on a sphere using FSL’s GPS tool (https://git.fmrib.ox.ac.uk/fsl/gps/-/tree/ master/). For each of the sampled directions, we calculated its dot-product with the original 90 directions (at bvals = 1000 s/mm^2^) and found its closest equivalent direction (i.e., maximum dot-product) from the actual 90 bvecs, resulting in a total of 32 single-shell volumes selected from the original multi-shell. Finally, we combined the selected 32 single-shell volumes with the bvals = 0 volumes to form the *LQ-LowAngular* dataset.

#### 2. LQ-LowSpatial

When scanning time is limited, it is possible to achieve better angular resolution by compromising on the spatial resolution for diffusion MRI. The *LQ-LowSpatial* dataset was designed to imitate this scenario, where the spatial resolution was compromised to achieve higher angular resolution. We downsampled the (minimally preprocessed) diffusion data from an isotropic spatial resolution of 1.25mm to 2mm while keeping the original shells and bvals/bvecs, resulting in a surrogate low-spatial-resolution dataset.

#### 3. LQ-LowAngular-LowSpatial

We also created the *LQ-LowAngular-LowSpatial* dataset, a surrogate low-angular and low-spatial resolution diffusion dataset, to reflect the more extreme poor data quality when advanced MRI imaging techniques are lacking (or time available for scanning is very short). This dataset was generated by downsampling the *LQ-LowAngular* dataset to an isotropic spatial resolution of 2mm.

#### 4. LQ-UKB

The UK Biobank diffusion MRI has a spatial resolution of 2×2×2 mm, which is closer to what is typically acquired on clinical scanners. Hence, we did not further degrade this spatial resolution. Instead, we only created its single-shell (low-angular-resolution) counterparts by extracting the b0 and bvals=1000 s/mm^2^ volumes, sampled at 32 diffusion directions.

### C. Structural and diffusion data post-processing

#### 1. Segmentation of thalamic masks

Thalamic masks for the left and right hemispheres were segmented for each subject in the HCP and UK Biobank datasets using FSL’s MIST (https://fsl.fmrib.ox.ac.uk/fsl/fslwiki/MIST), a sub-cortical segmentation tool that leverages complementary information from different MRI modalities to achieve accurate segmentation [60]. Although the HCP preprocessed data already included masks for the thalamus, these were not considered to be accurate enough and included too much white matter, as they were based on the T1 image only.

Three modalities were used for thalamus segmentation in MIST: T1, T2, and fractional anisotropy (FA), which was estimated using the diffusion tensor model [11] and FSL’s DTIFIT. To match the spatial resolution of the T1- and T2-weighted images, the FA images were upsampled from an isotropic resolution of 1.25 mm to 1 mm, while the T1- and T2-weighted images were downsampled from an isotropic resolution of 0.7 mm to 1 mm. The resulting thalamic masks were used as seed regions in tractography to construct white matter tracts projecting from the thalamus.

#### 2. Creation of other anatomical masks

We employed various regions-of-interest (ROIs) spanning the brain to create an augmented feature set that aims to robustly capture connectional characteristics of the Vim (even in low quality data). These ROIs included:

1. 75 cortical regions and 2 cerebellar parcels (one for cerebellar gray matter and the other for cerebellar white matter) per hemisphere, and the brainstem, segmented using Freesurfer [19, 20, 26, 27], released as part of the HCP preprocessed data.
2. 4 ROIs derived from the cerebellothalamic tract of the superior cerebellar peduncle (SCPCT) [57] warped into the individual subject space, which includes 3 white matter segments situated between the superior cerebellar peduncle (SCP) and the ipsilateral thalamus, as well as an additional parcel overlapping with the brainstem to account for the cerebellothalamic tract decussation. These ROIs are referred to as SCPCT-1, SCPCT-2, SCPCT-3, and SCPCT-brainstem.
3. 23 white matter segments extracted from the major white matter bundles projecting from or passing through the thalamus using the XTRACT atlas [62], including 6 segments from the Superior Thalamic Radiation (STR), 5 from the Acoustic Radiation (AR), 6 from the Anterior Thalamic Radiation (ATR), and 6 from the Optic Radiation (OR). These white matter ROIs were extracted in standard space and subsequently warped back into the individual space.
4. 5 white matter ROIs extracted from the fibre bundle joining thalamus and ipsilateral M1. This fibre bundle was created with thalamic voxels as the seed and M1 as both the waypoint and target mask, subsequently warped into standard space and averaged across subjects. These white matter ROIs were created in standard space and subsequently warped into individual subject space. These ROIs are referred to as M1-1, M1-2, M1-3, M1-4, and M1-5.
5. Another 5 white matter segments extracted from the fibre bundle joining thalamus and ipsilateral primary sensory cortex (S1). This fibre bundle was created similarly with thalamic voxels as the seed and S1 as both the waypoint and target mask, subsequently warped into standard space and averaged across subjects. These white matter ROIs were created in standard space and subsequently warped into individual subject space. These ROIs are referred to as S1-1, S1-2, S1-3, S1-4, and S1-5.

Table S1 provides a summary of the ROIs and details of how they were created.

#### 3. Fibre orientation estimation

Prior to tractography, fibre orientations were estimated for all diffusion MRI data by applying a parametric spherical deconvolution model using FSL’s BedpostX. In each voxel, up to three fibre orientations were estimated, along with their respective uncertainties [12, 13, 39, 55]. Two distinct deconvolution models were used to fit the crossing fibres in the multi-shell (*HQ-HCP*, *HQ-UKB*, and *LQ-LowSpatial*) and single-shell (*LQ-LowAngular*, *LQ-LowAngular-LowSpatial*, *LQ-UKB*) diffusion data. The former used a Ball-and-Sticks with zeppelins model [39, 55], while the latter used the standard Ball-and-Sticks model [12, 13]. The former approach models the diffusion coefficient using a Gamma distribution, enabling it to more effectively represent multi-shell diffusion MRI data. In contrast, the latter method models the diffusion coefficient with a single value, making it more appropriate for single-shell data.

#### 4. Tractography protocols

Probabilistic tractography was performed using FSL’s Probtrackx tool [12, 13, 35] in each dataset’s native individual T1 space. For the subjects that underwent repeat scans, the first-visit T1 scans served as the reference native space, to which the second-visit scans were registered. Since all the anatomical masks were in the (first-visit) T1 native space, an affine transformation matrix between the subject’s T1 and diffusion space was employed during fibre tracking, allowing the reconstructed streamline distributions to be directly sampled in the respective T1 space.

The anatomical masks included seeds (starting points of the streamlines), waypoints/targets (regions that streamlines must pass through to be valid), exclusion masks (regions that reject streamlines passing through them), and termination masks (regions that serve to stop streamlines running through them). Streamlines were seeded from the thalamic masks using a modified Euler integration with a step length of 0.5mm and 2000 steps per streamline, randomly initialised within a 1mm-radius sphere around each voxel centre. Streamlines were excluded if they entered the exclusion masks, or if the cosine of the angle between two consecutive steps exceeded 0.2.

A total of 5000 individual streamlines were drawn for each seed voxel and discarded if they reached the exclusion masks or did not meet the waypoint condition. The choice of waypoints/termination/exclusion masks varied depending on the target masks used in each run of tractography, with additional details given below. The output streamline distributions were corrected for the distance between the target and the seed mask, as the number of streamlines typically decreases with distance from the seed mask. A summary of the tractography options used in this study is provided in Table S2.

### D. Connectivity-driven approach

The connectivity-driven approach used in our study, reproducing the method of Akram et al. [1] and Bertino et al. [15], resulted in what we refer to as “connectivity-driven Vim” in the subsequent text. This approach identifies the Vim by finding the maximum probability of connection to M1 and contralateral dentate nucleus within the thalamic mask, generated via probabilistic tractography.

Streamlines seeded from the thalamus, targeting M1 and contralateral cerebellum, were generated via probabilistic tractography, yielding two “tract-density” feature maps for each target. These maps represent the white matter connectivity strength with their corresponding targets. To mitigate bias caused by differences in target volumes, the probability maps were normalised (setting the maximum to 1) and then thresholded to discard voxels characterised by low tract density (M1 at 50% of the tract density values within the thalamic mask; dentate at 15%). The thresholded M1 map was subsequently multiplied with the thresholded cerebellum map and binarised at a lower 30% threshold, resulting in the connectivity-driven Vim mask.

Note that the connectivity-driven approach was either applied to the high-quality data to generate HQ-Vim, serving as approximations of the ground truth location of the Vim, or to the low-quality data to produce low-quality connectivity-driven Vim. These low-quality connectivity-driven Vim segmentations were then compared with alternative approaches on low-quality data.

### E. Atlas-defined approach

The atlas-defined approach identifies the Vim based on the group-average location of the nucleus. To create a group-average Vim atlas, we transformed the binarised connectivity-driven Vim (derived from the high-quality data) to MNI152 1mm standard space [31] and averaged these across the training subjects to obtain a group-average Vim probability map. The transformation warp fields were obtained by non-linearly registering individual T1 to the same standard space. The voxel-wise probabilities in this Vim probability map represent the proportion of subjects that overlap at a given voxel. The group-average Vim probability map was subsequently warped back into individual T1 space and thresholded at 0.5 to define the “atlas-defined” Vim.

It is important to note that we only included a “reliable” subset of subjects when calculating the group-average Vim probability map. As the connectivity-driven approach has limited reliability even with high-quality data, we discarded subjects whose HQ-Vim were considered unreliable. An HQ-Vim must meet four criteria to be considered reliable:

1. Its volume exceeds 20 mm^3^. Given that the size of the Vim is approximately 4×4×6 mm [54], a resulting volume that is too small may suggest unreliable segmentation.
2. It contains only one connected component. This criterion aligns with the anatomical reality, given that the Vim structure is a single connected nucleus (in each hemisphere).
3. Its center-of-mass is located within 4 mm of the center-of-mass of the Vim cluster in the Thalamic DBS Connectivity Atlas [1], when transformed into native space.
4. Its correlation with the Thalamic DBS Connectivity Vim Atlas exceeds 0.5, again in native space.

Criteria 3) and 4) are used to exclude individual subjects’ HQ-Vim that deviate excessively from the atlas, whilst still allowing for a certain degree of individual variability. See Figure S1 for more details on the selection of the “reliable” subset.

### F. HQ-augmented approach

The goal of our approach is to leverage anatomical information in HQ data to infer the likelihood of a voxel belonging to the Vim, given a wide range of tract-density maps (multiple distinct tract bundles) derived from low-quality data as input features. The HQ-augmentation model was trained on the HCP dataset separately for each type of low-quality dataset. Using the HCP HQ data, we first generated the connectivity-driven Vim (referred to as HQ-Vim) as the “ground truth” location of the nucleus, serving as training labels in the model. Next, for each low-quality counterpart, we generated an extended set of tract-density features, targeting a wide range of ROIs, as the input features of the model. We hypothesise that the richer set of connectivity features serves to compensate for the primary tract-density features (connectivity with M1 and dentate), when those are compromised by insufficient spatial or angular resolution in low-quality diffusion MRI, thus making Vim identification less reliant on the tract-density features used in the connectivity-driven approach, and more robust to variations in data quality. During training, the model learns to use the extended set of connectivity features to identify the ground-truth Vim.

#### 1. Input features

The extended set of target ROIs comprised: 1) 75 ipsilateral cortical regions and the contralateral cerebellar white matter, derived from the Destrieux atlas [20]; 2) 23 white matter ROIs extracted from the XTRACT atlas [62]; 3) SCPCT-1, SCPCT-2, SCPCT-3, and SCPCT-brainstem, i.e., the 4 ROIs extracted from the cerebellothalamic tract of the superior cerebellar peduncle (SCPCT) [57], among which three are white matter segments lying between the thalamus and brainstem (SCPCT-1, SCPCT-2, and SCPCT-3), and one is overlapping with the brainstem (SCPCT-brainstem); 4) M1-1, M1-2, M1-3, M1-4, and M1-5, i.e., the white matter ROIs joining thalamus with ipsilateral M1; 5) S1-1, S1-2, S1-3, S1-4, and S1-5, i.e., the white matter ROIs joining thalamus with ipsilateral S1. These ROIs were chosen to span both grey matter and white matter regions. When data quality is compromised, tractography may fail to streamline into the grey matter, while it might be easier to reach the white matter segments, compensating the compromised gray-matter-related tract-density maps. The numbers refer to different locations of white matter segments along the corresponding fibre bundle.

For each target, we reconstructed streamlines with the thalamus as the seed mask and the selected target as a waypoint mask, producing a tract-density map within the thalamic voxels. The ipsilateral cerebellum and CSF were designated as exclusion masks, whilst the cortex served as termination regions. The streamlines were constrained to be ipsilateral, with the exception of the four SCPCT ROIs and the contralateral cerebellum serving as targets. Each of the resulting 113 tract-density maps was normalised by its maximum density value. Finally, six M1-related tract-density maps (targeting at M1 and five white matter segments from M1-1 to M1-5) were each multiplied by the following four tract-density maps (targeting the four SCPCT ROIs and the contralateral cerebellum) respectively, resulting in 6 *×* 5 product maps as additional connectivity features. These interaction features were designed to account for the intersection of the DTCp within the thalamic mask. The whole procedure resulted in a total of 143 features to be fed into the HQ-augmented model. Note that the input features were consistently located within the individual’s T1 space. This held true irrespective of the actual spatial resolution of the low-quality diffusion MRI. During the process of fibre tracking, the features were systematically resampled to align with the individual’s T1 space. This approach ensured uniform spatial correspondence across all dMRI datasets, irrespective of their original spatial resolution.

#### 2. Model setup and implementation

We used a Conditional Random Field (CRF) [63] to learn the mapping between the low-quality connectivity features (input features) and the HQ-Vim (training labels) using a regularised cross-entropy likelihood loss.

Formally, let **X** = [**x**_1_, **x**_2_, …**x***_V_]^T^* be a *V ×d* connectivity feature matrix for a given subject, where **x***_i_* is a *d ×* 1 vector representing the *d* connectivity features in voxel *i*, and **y** = [*y*_1_*, y*_2_*, …y_V_*]*^T^* is a *V ×* 1 vector containing the HQ-Vim labels. Suppose **T** is the one-hot encoding matrix of the HQ-Vim labels **y**, with *y_i_* mapped to a binary vector **t***_i_*, where *y_i_* = *k* corresponds to *t_ik_* = 1 and other elements in **t***_i_* are set to 0. Maximising the log-likelihood of reproducing HQ-Vim **y** given the low-quality connectivity features **X** is equivalent to minimising the following cross entropy (i.e., negative log likelihood):

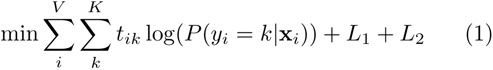

where *L*_1_ and *L*_2_ are penalty terms on the model parameters to prevent overfitting (these two terms are defined below).

The posterior probability *P* (*y_i_|***x***_i_*) is modelled by a CRF distribution. It consists of two terms: one models the likelihood of assigning a label to a given voxel without considering its neighbourhood, and the other accounts for the likelihood that neighbouring voxels have the same label. Mathematically:

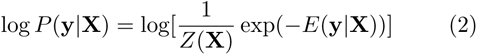

Here, *E*(**y***|***X**) models the cost of the label assignment **y** given the features **X**, and *Z*(**X**) is an image-dependent normalising term. *E*(**y***|***X**) is given as:

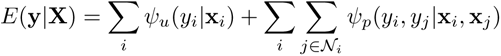

The unary cost *ψ_u_*(*y_i_|***x***_i_*) and pairwise cost *ψ_p_*(*y_i_, y_j_|***x***_i_,* **x***_j_*) measure the cost of assigning labels to voxels while encouraging similar labels for neighbouring voxels with similar features. Here *ψ_u_*(*y_i_|***x***_i_*) takes the form 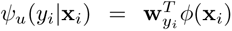, where *ϕ*(*·*) maps a feature vector **x***_i_* = [*x*_1_*, x*_2_*, …x_d_*] to a further expanded feature space in order to provide more flexibility for the parameterisation. **W** = [**w**_1_, **w**_2_] is the coefficient matrix to be learned from the data, each column containing the coefficients for the given class (i.e., belonging to the HQ-Vim or not). Here we chose a series of polynomials along with the group-average Vim probability (registered into native space) to expand the feature space. Specifically, *ϕ*(**x***_i_*) undergoes polynomial feature expansion, augmented by a group-average term. Each feature *x_k_* in **x***_i_* is transformed to include its corresponding polynomial terms 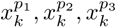, along with the original feature values. These are then concatenated with *g_i_*, the group-average Vim probability for the voxel, to form the expanded feature vector 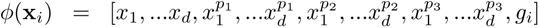. The powers of the polynomials *p*_1_ = 2*, p*_2_ = 0.5*, p*_3_ = 0.2 were chosen by testing a range of power values on an independent subset (see Figure S2), and *g_i_* is the group-average probability of voxel *i* classified as Vim (calculated across the training subjects).

The second pairwise cost encourages assigning similar labels to neighbouring voxels, particularly for those sharing similar connectivity features. We modelled this component as *ψ_p_*(**x***_i_,* **x***_j_*) = *ρµ*(*y_i_, y_j_*)*k* [*ϕ*(**x***_i_*)*, ϕ*(**x***_j_*)]. Here *k* [*ϕ*(**x***_i_*)*, ϕ*(**x***_j_*)] = exp(*−γ||ϕ*(**x***_i_*) *− ϕ*(**x***_j_*)*||*^2^) is a kernel function modelling the similarity between voxel *i* and *j* in the extended feature space, with length scale *γ*, chosen via cross-validation. *µ*(*·*) is a label compatibility function where *µ*(*y_i_, y_j_*) = 0 if *y_i_* = *y_j_* and *µ*(*y_i_, y_j_*) = 1 if *y_i_* ≠ *y_j_*.

Therefore, in a local neighbourhood, the kernel function penalises inconsistent label assignment of voxels that have similar features, thus allowing modelling local smoothness. *ρ* controls the relative strength of this pairwise cost weighted by *k*(*·*). Lastly, the *L*1 and *L*2 penalty enforced on **W** were included in the loss function (1) to prevent overfitting of the model. We used a mean-field algorithm to iteratively approximate the maximum posterior *P* (**y***|***X**) [63] summed across the subjects. The approximated posterior is maximised via gradient descent in a mini-batch form, where the connectivity feature matrix of each subject serves as a mini-batch, is demeaned and normalised, and sequentially fed into the optimisation problem. See supplementary materials for more details on model implementation.

### G. Evaluation of Vim localisation

The three approaches for Vim localisation on lowquality data were evaluated separately on the two subsets of subjects: the reliable subset of subjects, in which the HQ-Vim passed the four selection criteria, and the unreliable subset, in which the HQ-Vim did not meet the criteria. On the reliable subset, the corresponding HQ-Vim served as the ground truth (i.e., HCP HQ-Vim as ground truth for HCP; UKB HQ-Vim as ground truth for UKB); on the unreliable subset, the atlas-defined Vim served as the ground truth.

Two metrics, Dice coefficient and centroid displacement, were employed to assess the correspondence between the low-quality Vim localisation and the ground truth. The Dice coefficient measures the extent of overlap between two segmentations relative to their combined size. Specifically, for two segmentations, *A* and *B*, the Dice coefficient can be expressed as 2*|A ∩ B|/*(*|A|* + *|B|*), where *|A ∩ B|* represents the number of voxels classified as Vim in both *A* and *B*, whilst *|A|* + *|B|* represents the total number of Vim voxels in *A* and *B*. Since the HQ-augmentation approach and the atlas-based approach produce continuous Vim probability maps, calculating their Dice coefficients with the “ground truth” requires thresholding to binarise these maps *a priori*.

The other metric, centroid displacement, provides an alternative means of evaluating segmentation similarity that does not rely heavily on precise thresholding. It measures the Euclidean distance between the centroids of two Vim clusters, thus offering a measure of the spatial displacement between predicted and true Vim locations. This metric is also more clinically useful tha DICE in the context of DBS targetting. The centroid for the HQ-augmented Vim is calculated as the weighted average coordinates, where the weights correspond to the estimated posterior probability of a voxel being classified as Vim. A low threshold of 0.1 is applied to the output posterior probability map when calculating the centroid coordinate, which eliminates voxels that have a low likelihood of being classified as Vim. The centroid for the atlas-based Vim is computed in a similar fashion, using the group-average Vim probability as weights (once warped into native space) and also applying a 0.1 threshold. For the connectivity-driven Vim, the centroid coordinate is derived from the average coordinates of the binarised map.

## III. RESULTS

### A. Accuracy of the HQ-augmentation model on HCP surrogate low-quality data

As discussed above, the connectivity-driven approach may fail even on high-quality data. We therefore split the evaluation subjects into two subsets based on the reliability of the HCP HQ-Vim and made evaluations separately, against different ground truths (HQ-Vim for the reliable subset, atlas-driven for the other, as no reliable HQ-Vim exists for these subjects).

A total of 459 HCP subjects out of 1063 passed the four criteria and were selected as the reliable subset (see Figure 3 for the example contours of reliable HQ-Vim, and Figure 4C, 4F and 4I for contours of unreliable HQ-Vim). A group-average Vim probability map was calculated across these 459 subjects’ HQ-Vim (warped into standard space), and subsequently transformed back into the individual space and thresholded at 50% as the ground truth for the unreliable subset of HCP subjects. To further validate our choices of ground truth, we assessed the spatial proximity between this group-average Vim probability map and three sets of literature-based optimal stimulation points targeting the Vim for the treatment of tremor [22] in left hemisphere (as the three sets of coordinates were only provided for the left hemisphere). The group-average map overlapped with all three sets of optimal stimulation points (TABLE I). Notably, it overlapped with the peak location of an unweighted volume of tissue activated (VTA) frequency map with high probability.

**FIG. 2.**
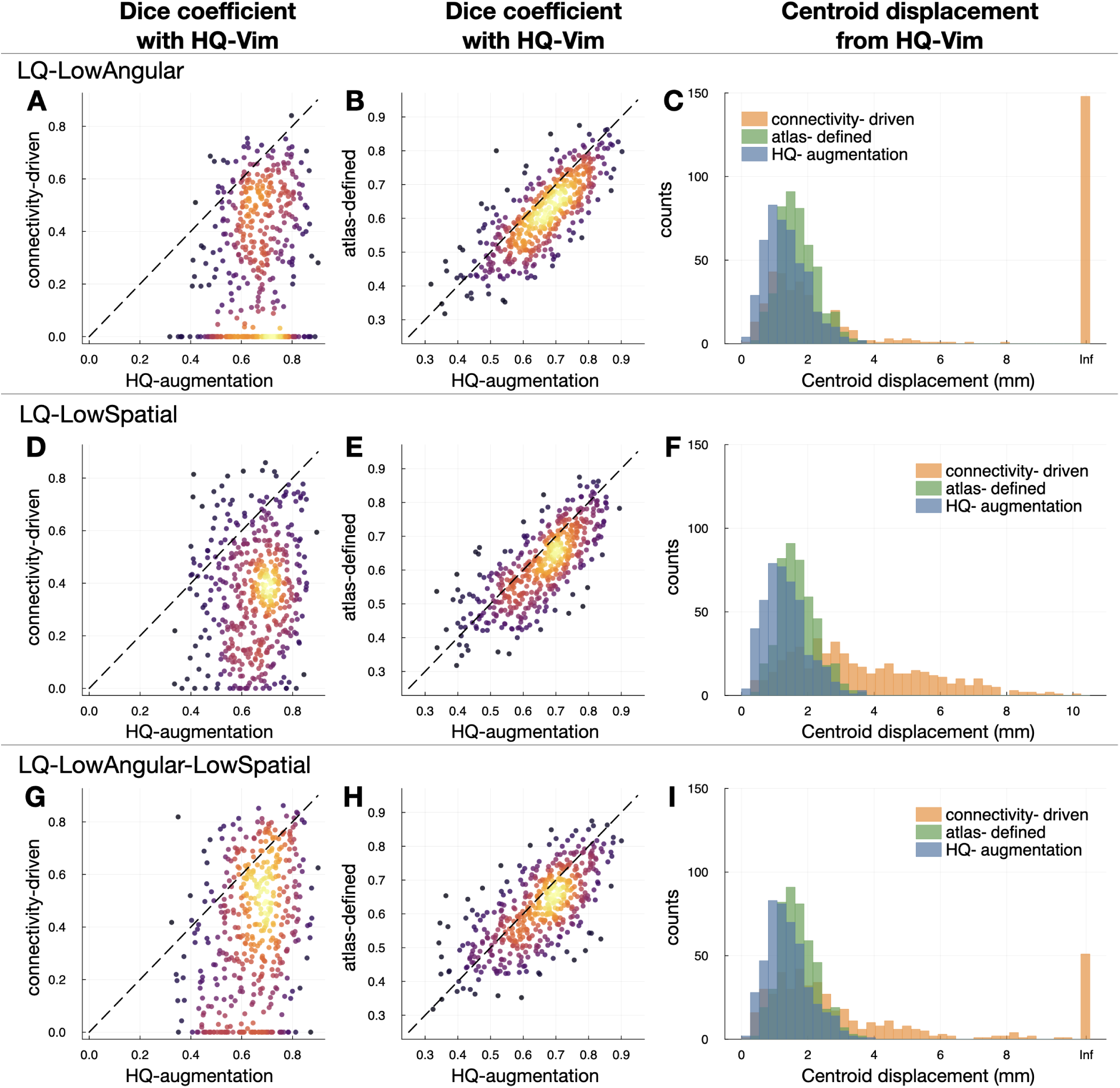
Accuracy of the HQ-augmentation model in reliable subjects. On each type of HCP surrogate low-quality data, we evaluated the HQ-augmentation, connectivity-driven, and atlas-defined approach against the HQ-Vim. **(A)** On *LQ-LowAngular* data, the HQ-augmentation model produced (in most subjects) a higher Dice coefficient (X-axis) with the respective HQ-Vim than the connectivity-driven approach (y-axis). Warmer colours are indicative of areas where data points are more densely populated. **(B)** On *LQ-LowAngular* data, the HQ-augmentation model also gave higher Dice coefficient with the respective HQ-Vim than the atlas-defined vim (y-axis). **(C)** On *LQ-LowAngular* data, the HQ-augmentation model (blue histogram) overall gave the smallest centroid displacement (from the HQ-Vim) than the connectivity-driven (orange) and atlas-defined (green) approach. **(C), (D) and (F)** Equivalent plots of on *LQ-LowSpatial*. **(G), (H) and (I)** Equivalent plots on *LQ-LowAngular-LowSpatial*. For a more direct representation of the summary statistics, please refer to Figure S3 which displays boxplots of the Dice coefficients.

**FIG. 3.**
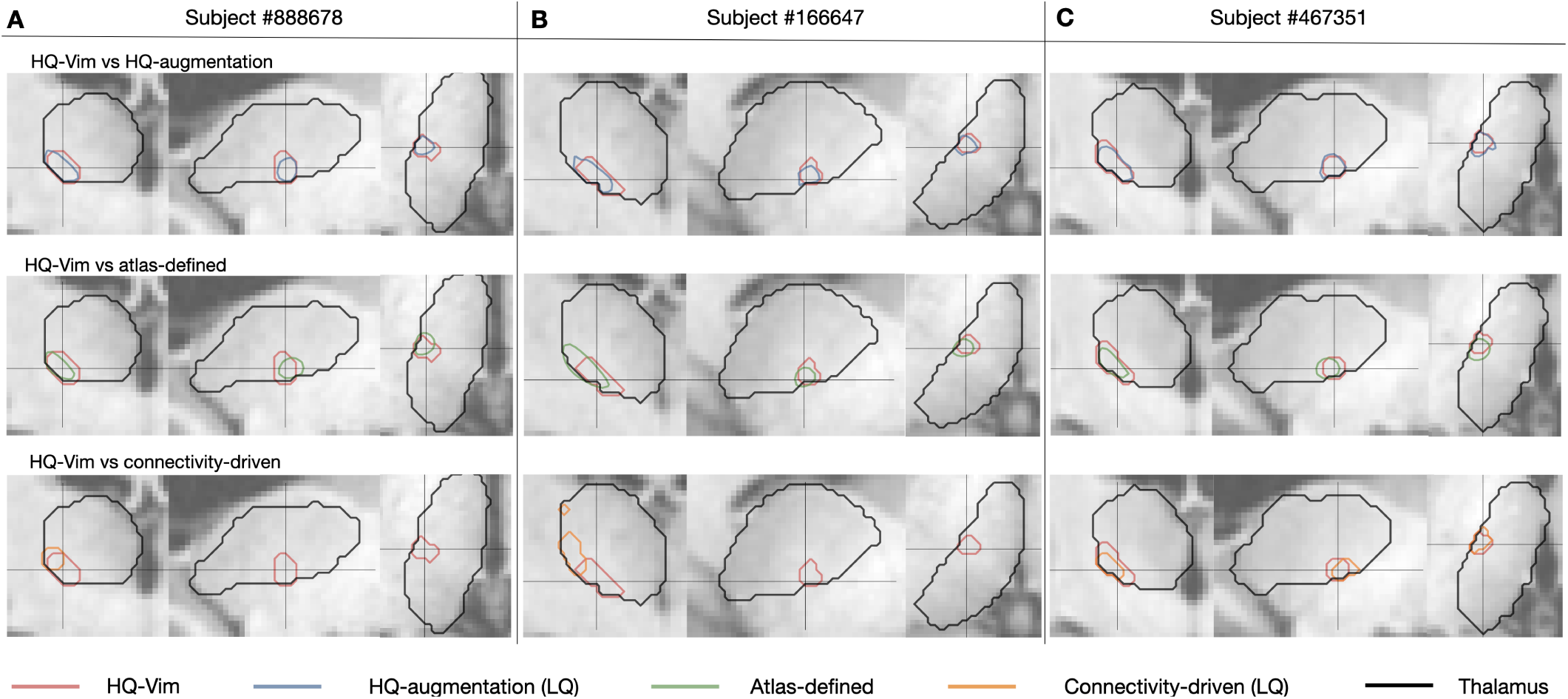
Example contours of HQ-Vim (ground truth), overlayed with the HQ-augmented or connectivity-driven Vim from HCP low-quality data, or by the atlas-defined Vim. **(A)** Contours of HQ-vim (red) versus *LQ-LowAngular* Vim (blue: HQ-augmentation; orange: connectivity-driven), or the atlas-defined Vim (green), of an example subject. **(B), (C)** are similar to (A) but for two other example subjects.

**FIG. 4.**
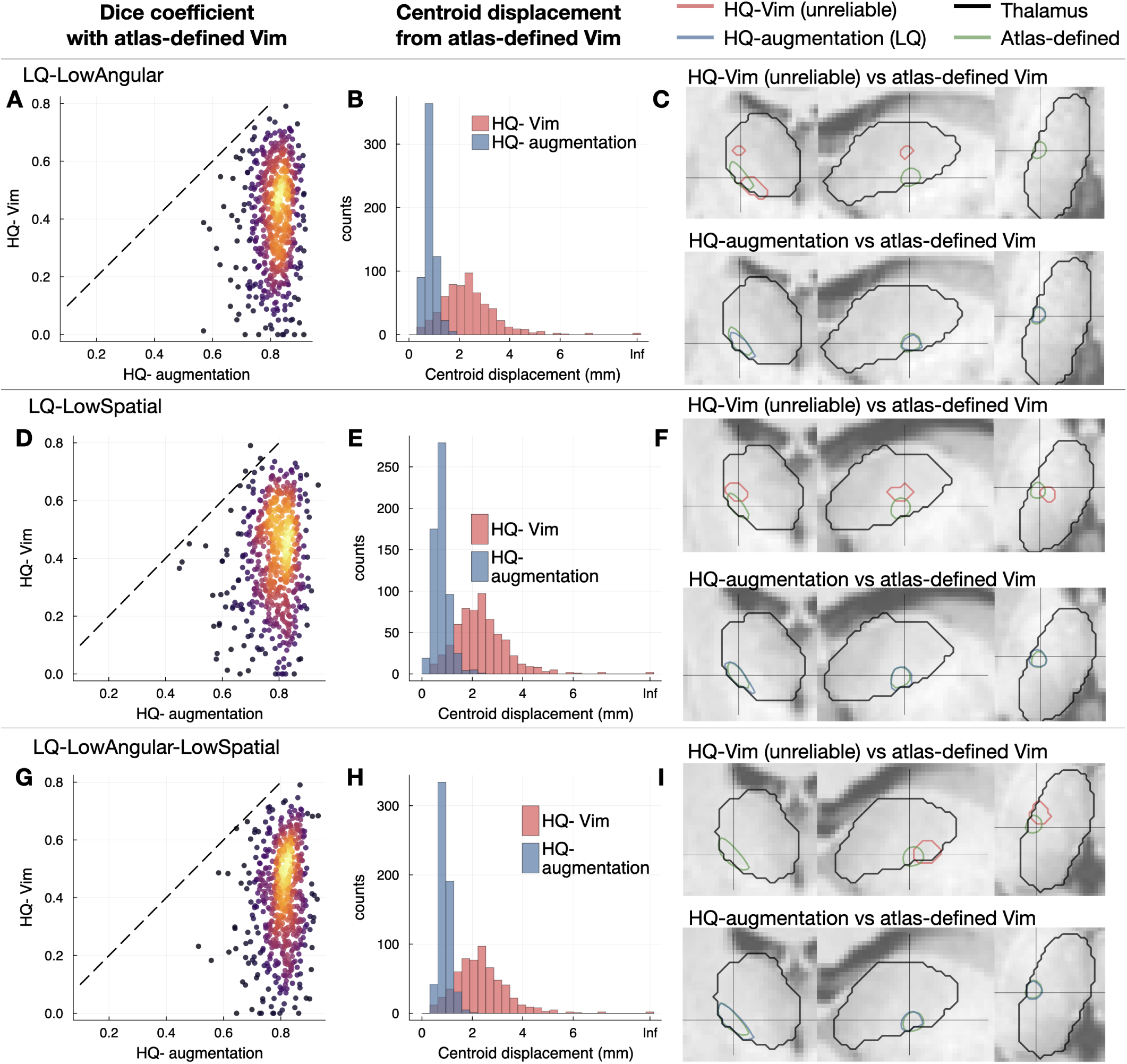
Accuracy of HQ-augmentation in unreliable subjects. For the left-out subjects in which HQ-Vim is untrustworthy, the atlas-defined Vim served as “ground truth” instead. **(A) and (B)** The HQ-augmentation approach (on *LQ-LowAngular*) gave higher Dice coefficient and smaller centroid displacement with the atlas-defined Vim even than the HQ-Vim. Warmer colours are indicative of areas where data points are more densely populated. **(C)** Example contours of HQ-Vim (unreliable), atlas-defined Vim (green) and HQ-augmented Vim (blue). The HQ-augmentation approach on *LQ-LowAngular* produced Vim closer to the group-average location of Vim. **(D), (E) and (F)** Equivalent plots of **(A), (B) and (C)**, on *LQ-LowSpatial* data. **(G), (H) and (I)** Equivalent plots of **(A), (B) and (C)**, on *LQ-LowAngular-LowSpatial* data.

**TABLE I.** Spatial proximity between the group-average Vim probability map and the literature-based optimal stimulation points (left hemisphere). The group-average Vim probability map calculated from the reliable subset overlapped with three sets of literature-based optimal stimulation points. The overlap probability represents the proportion of subjects whose Vim mask contains the given target location. The classical target center of gravity was defined by Horn et al. based on literature-reported effective contact location. The other two sets were defined by Elias et al. 2021 on the basis of volume of tissue activated (VTA) stimulation maps, either weighted by the corresponding VTA percentage change from baseline (clinically-weighted hotspot coordinates), or unweighted but thresholded at top 10% (unweighted frequency map peak coordinates).

| Literature-based stimulation points | I: classical target center of gravity | II: clinically-weighted hotspot | III: unweighted frequency map peak |
| --- | --- | --- | --- |
| MNI coordinates (x, y, z) | (-13.1, -19.3, -2.1) | (-17.3, -13.9, 4.2) | (-16.5, -17.0, 1.2) |
| Overlap probability (HCP) | 0.51 | 0.37 | 0.96 |
| Overlap probability (UKB) | 0.62 | 0.23 | 0.95 |

Next, we compared three Vim localisation approaches against the “ground truth” for each type of HCP surrogate low-quality data: 1) the atlas-defined Vim; 2) the connectivity-driven approach, 3) the HQ-augmented Vim. When evaluated against the respective HQ-Vim, the HQ-augmented Vim outperformed the atlas-defined Vim and the low-quality connectivity-driven Vim for all three types of HCP low-quality data (Figure 2), producing higher Dice coefficient with the respective HQ-Vim and smaller Euclidean distance from the HQ-Vim’s center-of-mass. In particular, on *LQ-LowAngular* (Figure 2A, 2B and 2C) and *LQ-LowAngular-LowSpatial* data (Figure 2G, 2H and 2I), the connectivity-driven approach often failed to generate any clusters due to the degraded angular resolution, resulting in “zero” identified Vim volume and thus having zero Dice coefficient with the HQ-Vim. In this case, its Euclidean distance from the HQ-Vim’s center-of-mass was set to Infinity. In contrast, the HQ-augmentation model invariably succeeded in finding a coherent Vim cluster that was close to the respective HQ-Vim, suggesting that the HQ-augmentation model is more capable of reproducing Vim localisation than the connectivity-driven approach when high-quality data is unavailable. Furthermore, the HQ-Vim approach produced higher Dice coefficients and smaller centroid displacements with the HQ-Vim than the atlas-defined approach, suggesting that the HQ-augmented Vim preserved more inter-individual variations of the Vim than a fixed standardised atlas. Obviously, the subset of HQ-Vim as evaluation “ground truth” was defined to resemble the standardised Vim atlas (i.e., excluding those having a large mismatch with the atlas), which thus limits inter-individual anatomical variability in the first place. It is notable that the HQ-augmentation approach produced results that were closer to the HQ-vim than the atlas-defined Vim, even within this limited margin of anatomical variability (see Figure 3 for contours of reliable HQ-Vim, atlas-defined, HQ-augmented and connectivity-driven Vim).

On the subset of subjects for which the HQ-Vim were regarded as unreliable, we evaluated the accuracy of the HQ-augmented Vim against the atlas-defined Vim (Figure 4). Although the atlas-defined Vim cannot fully account for individual variability, it may still serve as a version of the “gold standard”, as the unknown ground truth location of the Vim is not expected to be far from the atlas position. For all three types of low-quality data, the HQ-augmentation approach on the low-quality data gave higher Dice coefficient and smaller centroid displacement with the atlas-defined Vim, even outperforming the connectivity-driven approach applied on high-quality HCP data. This suggests that the HQ-augmentation approach is more robust to corruptions of data quality. Similarly, this evaluation (on unreliable subjects) is unavoidably somewhat circular, in the sense that these subjects are defined by a large mismatch between HQ-Vim and atlas-based Vim, but it is nevertheless notable that the HQ-augmentation results (which are driven by tractography) are closer to the atlas results than the HQ-Vim (see Figure 4C, 4F and 4I for example contours of unreliable HQ-Vim, the HQ-augmented Vim on each low-quality dataset, and the atlas-defined Vim).

Overall, the above results demonstrate that the HQ-augmentation model is superior to existing approaches in its robustness to data quality, as well as its ability to preserve individual variability.

### B. Generalising the HQ-augmentation model to the UKB surrogate low-quality dataset

One limitation of our evaluations so far is that both training and test data were derived from the HCP dataset (albeit corrupted with downsampling). To be useful, it is imperative for our model to be generalisable to unseen protocols, as collecting large datasets for training purposes in clinical contexts may be impractical. Here we leveraged the UKB diffusion MRI dataset to test the generalisability of the HQ-augmentation model trained on HCP. As described in the methods section, we created a *LQ-UKB* dataset by degrading the original 3T UKB dataset, and next applied the HQ-augmentation model trained on *LQ-LowAngular-LowSpatial* to its connectivity features, producing a predicted Vim probability map for each UKB subject.

The evaluations on the UKB dataset were similar to those on HCP, where the evaluation subjects were divided into a reliable subset with UKB HQ-Vim as ground truth and an unreliable subset with atlas-defined Vim as ground truth. The UKB HQ-Vim were derived from the original multi-shell UKB dataset via the connectivity-driven approach. The UKB atlas-defined Vim was derived from the UKB version of group-average Vim probability map, created by averaging the reliable UKB HQ-Vim, as it is more representative of the age population in UKB than the HCP group-average Vim probability map. The selection criteria were the same as HCP (Figure S1B). The group-average Vim probability map calculated from the subset of reliable HQ-Vim, though slightly different from the one calculated from the HCP subjects, also overlappped with the literature-based optimal stimulation points for tremor (see TABLE I).

Similarly to HCP, the performance of the HQ-augmentation model was compared with the connectivity-driven Vim on LQ-UKB and the atlas-defined Vim. Although purely trained on HCP low-quality features, the HQ-augmentation model again produced higher Dice coefficient and smaller centroid displacement than the connectivity-driven and atlas-defined approach (Figure 5A) with the UKB HQ-Vim, suggesting its robustness to variations in data acquisition as well as its ability to capture individual variability. Furthermore, when evaluated against the atlas-defined Vim, this HCP-transferred HQ-augmentation model generated Vim estimates that were much closer to the atlas than the UKB HQ-Vim, indicating again that the HQ-augmentation approach, even trained on a different dataset, was even more reliable than the connectivity-driven approach applied to high-quality data (Figure 5B). Overall, these results demonstrate that the HQ-augmentation approach trained on HCP data can indeed generalise to other lower-quality datasets acquired using different protocols. A possible reason for this generalisability is that the HQ-augmentation model trained on HCP has already learned some general features and patterns that can be useful for predicting Vim on UKB data; hence, leveraging the anatomical knowledge gained from HCP improves its performance on another related dataset, e.g., LQ-UKB. The HCP dataset itself contains a diverse range of subjects, and thus the model has learned to handle the variability in the LQ-UKB to some extent.

**FIG. 5.**
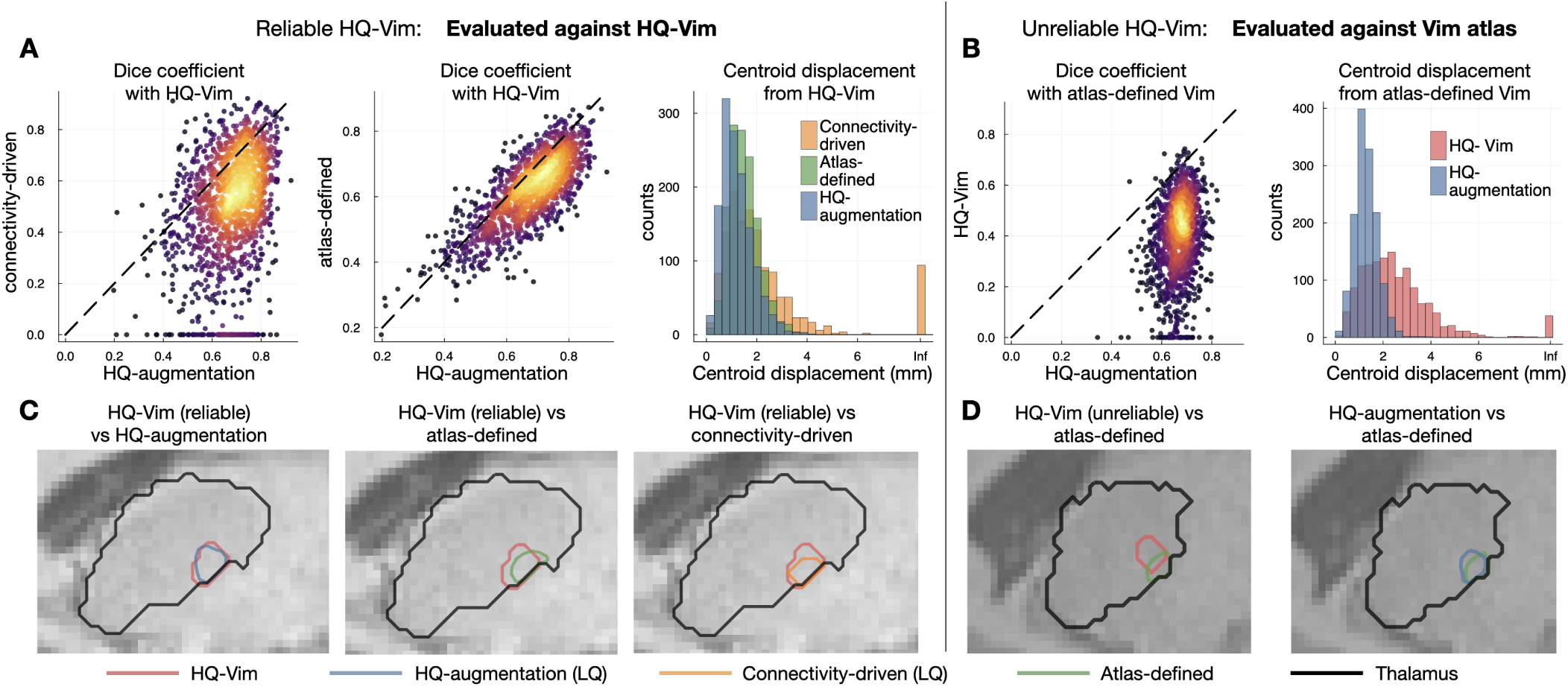
Generalisability of HQ-augmentation to UKB. The HQ-augmentation model trained on HCP was applied to the UKB low-quality features (blue) as HQ-augmented Vim. **(A)**. When using the UKB HQ-Vim as ground truth, the HQ-augmentated Vim has higher Dice coefficient and smaller centroid displacement with the UKB HQ-Vim, than the atlas-defined Vim (green) and the connectivity-driven Vim using low-quality features (orange). **(B)** When using the atlas-defined Vim as ground truth, the HQ-augmentation model using low-quality features even gave more reliable Vim than the UKB HQ-Vim (red), which used HQ features to target Vim. **(C)** and **(D)**. Contours of Vim, identified by each approach.

### C. Reliability analysis of the HQ-augmentation approach on HCP and UKB

We conducted a comprehensive reliability analysis for the three approaches at the individual level to assess their consistency in producing Vim localisation despite variations in data quality and across scanning sessions. It is crucial for a localisation approach to produce consistent results not only across different scanning sessions but also across varying data quality, as its clinical value lies in capturing the true underlying anatomical variability unique to each individual, rather than reflecting variabilities arising from noise or methodological factors. Our reliability assessment consisted of two dimensions: 1) consistency of Vim localisations across scans with different quality levels for a given subject (referred to as across-quality reliability); and 2) correspondence of Vim localisations across different scanning sessions of the same quality level for a given subject (referred to as test-retest reliability). Prior to conducting the reliability analysis, we trained an HQ-augmentation model specifically for the high-quality data, using the HQ-Vim as training labels and the extended set of connectivity profiles derived from high-quality data instead as input features. The HQ-augmented model tailored to high-quality data was applied to the left-out subjects to produce a “high-quality version” of HQ-augmented Vim per subject. This version was then compared with the HQ-augmentation outputs from low-quality datasets (to measure its across-quality reliability) or retest sessions (to measure its test-retest reliability).

To assess across-quality consistency of the HQ-augmentation approach, we calculated the Dice coefficient and centroid displacement between the HQ-augmented Vim derived from each low-quality dataset (*LQ-LowAngular*, *LQ-LowSpatial*, *LQ-LowAngular-LowSpatial*) and its high-quality counterpart (i.e, the high-quality version of HQ-augmented Vim) per subject, resulting in six measurements per subject (2 metrics *×* three pairs of low-versus-high comparisons). Similarly, we also assessed the across-quality consistency of the connectivity-driven approach, by comparing the connectivity-driven Vim derived from each low quality dataset against the respective HQ-Vim, again resulting in six measurements. On both UKB and HCP, the HQ-augmentation approach produced substantially more consistent Vim localisation results than the connectivity-driven approach across datasets of different quality (Figure 6). Both approaches had compromised “across-quality” consistency on the unreliable subset, demonstrated by the smaller Dice coefficient and higher centroid displacement (Figure 6, light colours).

**FIG. 6.**
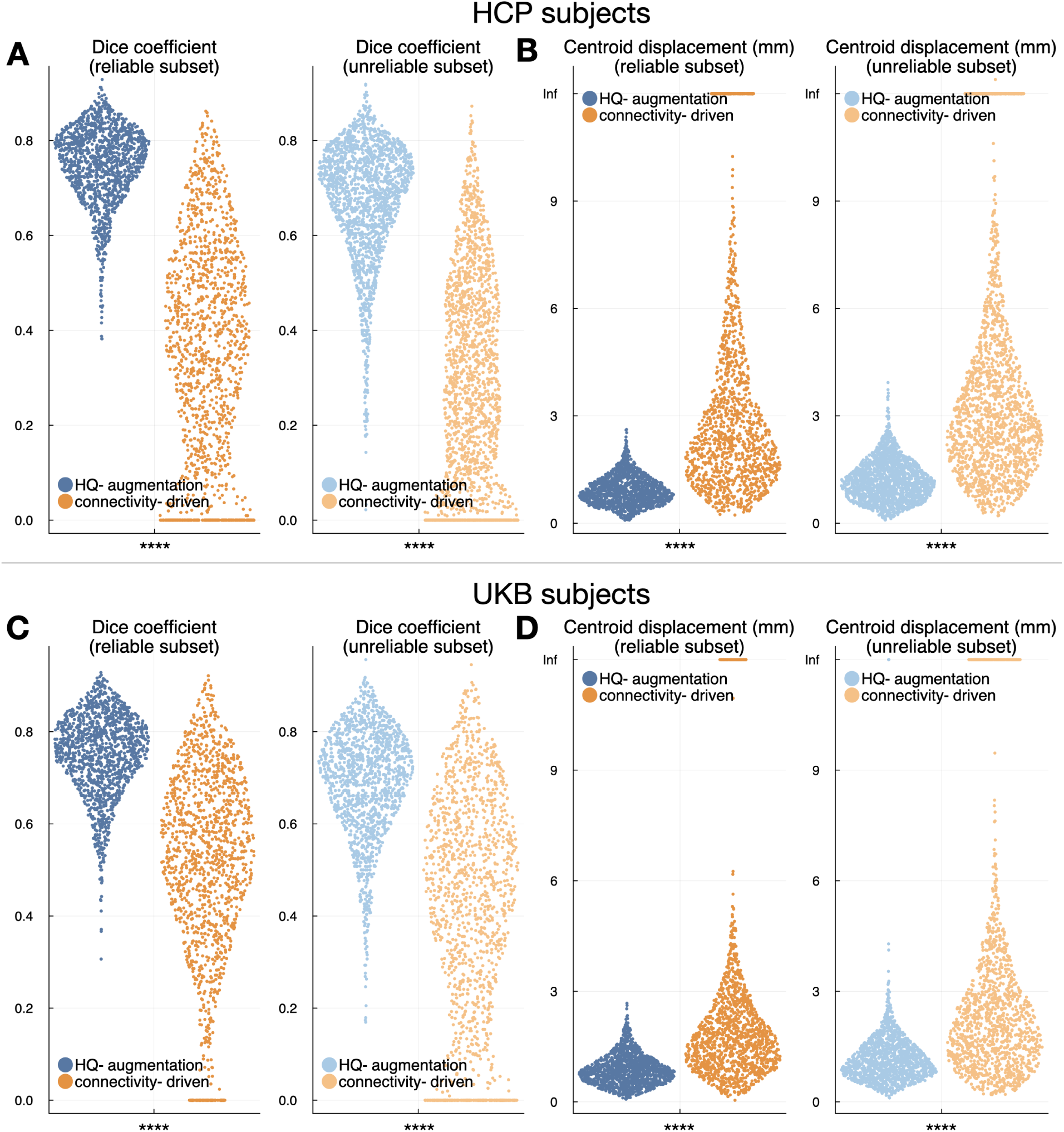
Across-quality consistency of the HQ-augmentation and connectivity-driven approach. The HQ-augmentation approach is significantly more consistent than the connectivity-driven approach (paired t-tests, Bonferroni-corrected by 8 tests in total), producing higher Dice coefficient and smaller centroid displacement between the outputs derived from different data quality, both across the reliable subset (dark blue and orange) and unreliable subset (light blue and orange). **(A)** Blue: Dice coefficient between the HQ-augmentation results on high-quality data (HQ-HCP) and the low-quality data (*LQ-LowAngular*, *LQ-LowSpatial*, *LQ-LowAngular-LowSpatial*), pooled together (i.e., each dot represents a single high-vs-low comparison); orange: equivalent plots for the connectivity-driven approach. **(B)** Equivalent plots to (A), using centroid displacement as the consistency metric. **(C)** Dice coefficient between the HQ-augmentation results on HQ-UKB and LQ-UKB (blue) and between the connectivity-driven results on HQ-UKB and LQ-UKB (orange). **(D)** Equivalent plots to (C), using centroid displacement as the consistency metric. Dark colours indicate reliable subjects (i.e., their HQ-Vim are trustworthy), while light colours indicate the opposite. Statistical significance is denoted as follows: **** for p<0.0001.

The test-retest reliability of the two approaches displayed a similar pattern. To assess the test-retest reliability of the HQ-augmentation approach, we applied the HQ-augmentation model trained on the multi-shell first-visit data to the second-visit connectivity features, producing the HQ-augmented Vim for the repeat sessions, and calculated the Dice coefficient and centroid displacement between first-visit and second-visit HQ-augmented Vim. The test-retest reliability of the connectivity-driven approach was likewise measured by the similarity between the connectivity-driven Vim derived from the first-visit and second-visit M1/dentate connectivity features respectively. The HQ-augmentation approach outperforms the connectivity-driven approach in providing substantially higher Dice coefficient and smaller centroid displacement between sessions (Figure 7). Similarly, both approaches had compromised test-retest reliability on the unreliable subset (Figure 7, light colours).

**FIG. 7.**
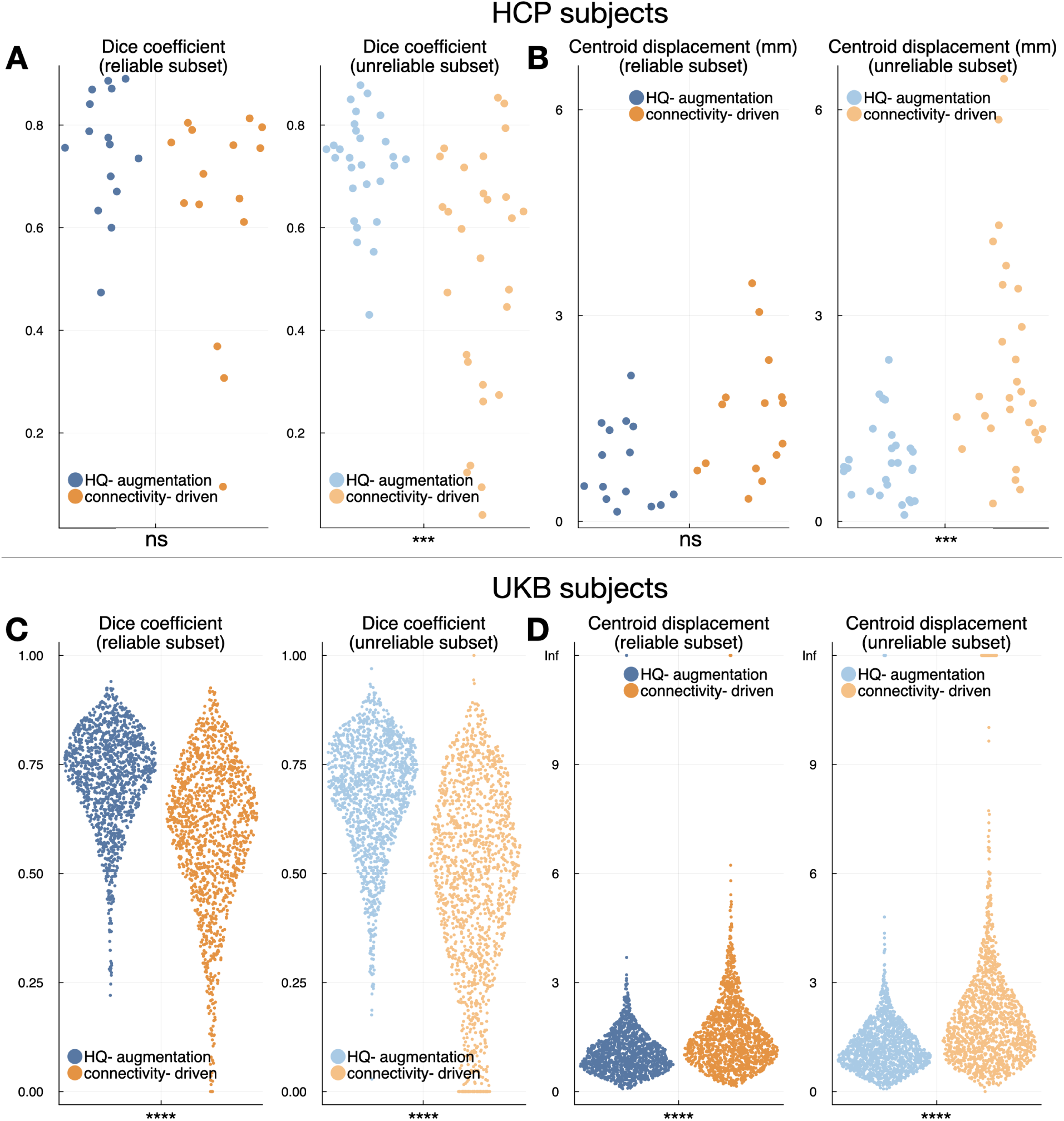
Test-retest reliability of the HQ-augmentation and connectivity-driven approaches. The HQ-augmentation approach is significantly more consistent than the connectivity-driven approach (8 paired t-tests, Bonferroni-corrected by 8 tests in total) across scanning sessions, producing higher Dice coefficient and smaller centroid displacement between the outputs derived from the first-visit and repeat scans, both across the reliable subset (dark blue and orange) and unreliable subset (light blue and orange). **(A)** Blue: Dice coefficient between the HQ-augmentation outputs on first-visit and repeat scans across 43 HCP subjects that had received retest sessions; orange: equivalent plots for the connectivity-driven approach. **(B)** Equivalent plots to (A), using centroid displacement as the consistency metric. **(C)** and **(D)** Equivalent plots to (A) and (B) for 2,760 UKB subjects, using centroid displacement as the consistency metric. Dark colours indicate reliable subjects (i.e., their HQ-Vim are trustworthy), while light colours indicate the opposite. Statistical significance is denoted as follows: ns for non significant, *** for p<0.001, and **** for p<0.0001.

## IV. DISCUSSION

We proposed an Image Quality Transfer (IQT) approach, HQ-augmentation, for improving the accuracy of localising the ventral intermediate nucleus of the thalamus (Vim), a common target for deep brain stimulation and MR-guided ultrasound. This approach leverages anatomical information (i.e., approximate ground truth location of the nucleus) obtained from high-quality HCP to augment Vim identification on the low-quality data, by learning to produce Vim localisation as close as possible to the high-quality “ground truth” using a wide range of white matter connectivity features derived from the low-quality data. Our results demonstrate that the HQ-augmentation approach is more accurate in estimating Vim, and more robust to data corruptions, than its predecessor, the connectivity-driven approach, especially when high-quality data is unavailable; moreover, it proves more reliable than the connectivity-driven approach even when the latter uses high-quality information to localise Vim. The HQ-augmentation approach performs consistently well across different data qualities and scanning sessions, surpassing the performance of the connectivity-driven approach. Furthermore, the HQ-augmentation approach preserves more inter-individual anatomical variability than the atlas-based approach. Importantly, our HQ-augmentation model trained on HCP surrogate low-quality data is generalisable to UKB low-quality data, outperforming the connectivity-driven approach in accuracy and reliability while preserving the anatomical variability unique to UKB individual subjects. Being a reliable and generalisable tool, our approach is valuable in clinical contexts where large samples of data collection and retraining are impractical.

Why is the HQ-augmentation approach capable of reproducing HQ-Vim on low-quality data, while the connectivity-driven approach using M1 and dentate tract-density features struggles to do so in isolation? The HQ-augmentation model leverages connectivity features with other target ROIs, which may also contain information about the anatomical location of Vim. For instance, Vim is anterior to thalamic clusters exhibiting high diffusion connectivity to S1, the primary sensory area. Consequently, tract-density with S1 serves as a “negative” feature, providing evidence against the likelihood of a given voxel being classified as Vim. Connectivity targeting white matter segments can also be informative, particularly when diffusion data has limited angular resolution. This is because middle point targets are more accessible for streamlines to reach compared to more distant cortical ROIs. For example, streamlines originating from a seed thalamic voxel might not reach M1 but could still reach white matter segments between the ipsilateral M1 and the thalamus. In such cases, tract-density features with these white matter ROIs may compensate for the inadequate M1 connectivity feature, resulting in Vim segmentation that remains close to the one defined by high-quality M1 and dentate tract-density features. In essence, the HQ-augmentation model synthesises evidence from an extended set of connectivity features to determine the most probable location of the “true” Vim.

When primary connectivity features defining Vim fail to provide reliable identification due to limited data quality, the remaining connectivity features may still guide the model to produce Vim segmentation close to its highquality counterpart. This makes the model more robust against data quality degradation. It is worth noting that the HQ-augmentation model generates Vim segmentation closer to the atlas than the HQ-Vim, which is defined by high-quality M1 and dentate tract-density features. This is due to the incorporation of a “prior” feature representing the group-average location of Vim. The HQ-augmentation approach may converge to the baseline, i.e., the atlas-defined Vim, when the entire extended set of connectivity features is too noisy to provide reliable Vim identification.

The proposed HQ-augmentation model falls within the broader category of Image Quality Transfer (IQT), a set of techniques aimed at propagating information from rare high-quality data to more prevalent low-quality datasets. Usually, this transfer entails enhancing the image resolution or information content, facilitating subsequent analyses. IQT typically relies on matched pairs of high- and low-quality data to establish a mapping from low-quality features to the corresponding high-quality information [2, 3, 25]. Although closely related to these previous IQT techniques, our HQ-augmentation approach differs slightly as it does not primarily focus on enhancing lower-quality images to augment subsequent analysis. Instead, it directly learns the mapping between high-quality Vim labels and lower-quality connectivity features. Once learned and applied to unseen image sets, the mappings “augment” the low-quality connectivity features to produce predicted Vim probability maps as if they were derived from their high-quality counterparts. An alternative option for augmenting Vim localisation on low-quality data could involve enhancing the low-quality image content via IQT first, followed by identifying Vim using the connectivity-driven approach on the enhanced image. However, this route was not chosen because the connectivity-driven approach can fail even on high-quality data. Therefore, in addition to high-quality image transfer, it is equally important to develop an approach that is more reliable and flexible than existing methods, even when high-quality data is available. Nevertheless, it is worth exploring whether enhancing low-quality image content via IQT first, followed by the HQ-augmentation model, might further improve Vim localisation on low-quality data.

The HQ-augmentation approach has several limitations. First, it relies on the availability of high-quality data to train the model, which may not always be accessible in certain research or clinical settings. Although our model demonstrates generalisability from HCP to UKB datasets, it is important to assess its performance on other lower-quality datasets and populations to ensure broad applicability. Moreover, given the variation in MRI acquisition protocols and hardware, the model may require retraining or fine-tuning for specific clinical or research contexts. Second, the current model is based on a machine learning approach, which relies on userdefined features and representations of the data. Recent advancements in deep learning, specifically in the domain of medical image analysis, may offer alternative solutions with improved performance and adaptability to various imaging contexts. Investigating the integration of deep learning methods, such as convolutional neural networks (CNNs), with the HQ-augmentation framework, could lead to further advancements in deep nuclei segmentation on low-quality data. Lastly, while the HQ-augmentation approach demonstrates improved performance compared to existing techniques, there is still room for refinement. Exploring additional image features, optimising the model architecture, and incorporating advanced regularisation techniques may further improve the segmentation accuracy and robustness of the model, ultimately benefiting clinical decision-making and personalised patient care.

Finally, we note that our approach is not exclusively applicable to the Vim nucleus. It can be generalised to any brain region for which detailed connectivity information is available (for example from animal models). This adaptability is particularly suitable for subcortical regions, in cases where the connectivity in animal models, such as macaques (where we can use tracers), is similar to that of humans.

## V. CODE AVAILABILITY

The source code for the model and the data analysis procedures are publicly accessible. They are hosted on the FMRIB Git repository and can be accessed via the following URL:

- Julia implementation: https://git.fmrib.ox.ac.uk/yqzheng1/hqaugmentation.jl;
- Python implementation: https://git.fmrib.ox.ac.uk/yqzheng1/python-localise.

## VI. ACKNOWLEDGEMENTS

SMS is supported by the Wellcome Trust Collaborative Award (215573/Z/19/Z), and MRC Mental Health Pathfinder grant (MC_PC_17215). SJ is supported by a Wellcome Senior Fellowship (221933/Z/20/Z) and Wellcome Collaborative Award (215573/Z/19/Z). The Wellcome Centre for Integrative Neuroimaging is supported by core funding from the Wellcome Trust (203139/Z/16/Z). The computational aspects of this research were partly carried out at Oxford Biomedical Research Computing (BMRC), which is funded by the NIHR Oxford BRC with additional support from the Wellcome Trust Core Award Grant Number 203141/Z/16/Z. We are grateful to UK Biobank for making this resource available (data access application 8107), and to the UK Biobank participants for dedicating their time to make this data possible. We are also grateful to the Human Connectome Project, WU-Minn Consortium (Principal Investigators: David Van Essen and Kamil Ugurbil; 1U54MH091657) funded by the 16 NIH Institutes and Centers that support the NIH Blueprint for Neuroscience Research; and by the McDonnell Center for Systems Neuroscience at Washington University.

## Supplemental Materials

### SELECTION CRITERIA OF RELIABLE HQ-VIM

The subjects were split into two subsets, depending on the reliability of HQ-Vim. A subject’s HQ-Vim has to pass 4 criteria in order to be accepted as “reliable”:

1. the HQ-Vim’s volume exceeds 20mm^3^;
2. the HQ-Vim contains one blob;
3. Its correlation with the Vim from Thalamic DBS Connecitivity Atlas [1] is larger than 0.5;
4. Its center-of-mass is within 4mm of the center of mass of the Thalamic DBS Connecitivity Dentate Atlas;

The thresholds in 3. and 4. are shown as vertical lines in Figure S1. 459 out of 1063 HCP subjects and 1445 out of 2760 UKB subjects passed all four criteria. These selection criteria exclude subjects whose HQ-Vim clusters lie too far away from the atlas to be considered as trustworthy, while preserving the inter-individual anatomical variability of the structure as much as possible. Note that, however, passing all the selection criteria does not necessarily guarantee the selected HQ-Vim as the perfect “ground truth”. Instead, this only suggests that the HQ-Vim may serve as the ground truth Vim with relatively high confidence, as it is the best available estimate of the ground truth location of the Vim.

**FIG. S1.**
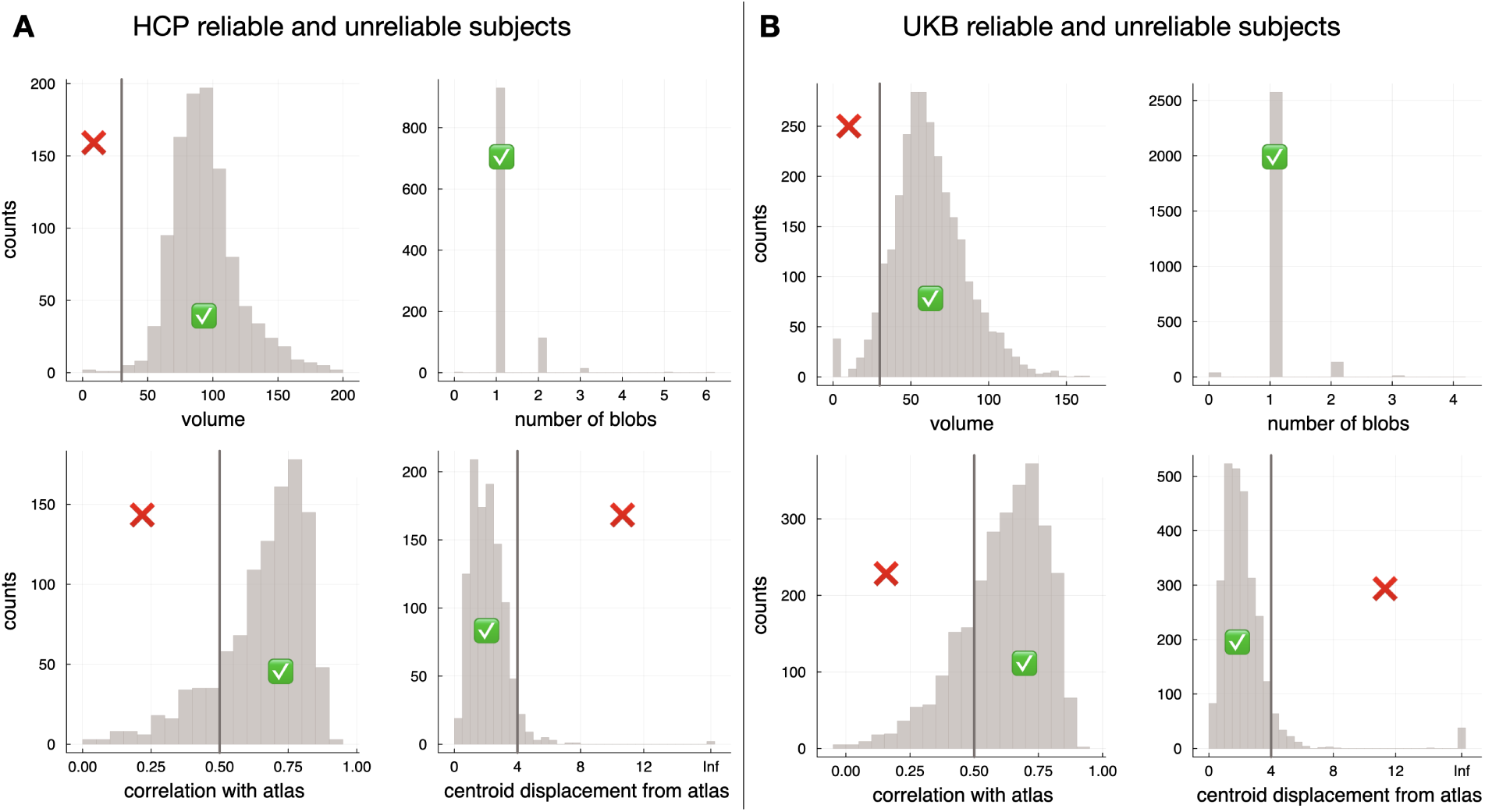
Split between the reliable and unreliable subsets. **(A)** Histograms of HQ-Vim’s volume (mm^3^), number of blobs, correlations and centroid displacement with the Thalamic DBS Connecitivity Atlas, for HCP subjects. **(B)** Equivalent plots to (A), for UKB subjects. Ticks indicate that the subjects pass the respective criteria, while crosses suggest their HQ-Vim were rejected as untrustworthy.

### THE POWER OF THE POLYNOMIALS

This section describes how we chose the power of polynomials to expand the feature space 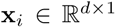. Specifically, *ϕ*(**x***_i_*) undergoes polynomial feature expansion, augmented by a group-average term. Each feature *x_k_* in **x***_i_* is transformed to include its corresponding polynomial terms 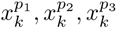, along with the original feature values. These are then concatenated with *g_i_*, the group-average Vim probability for the voxel, to form the expanded feature vector 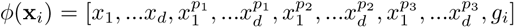. The powers of the polynomials *p*_1_ = 2*, p*_2_ = 0.5*, p*_3_ = 0.2 were chosen by testing a range of power values on an independent subset (Figure S2), which appeared to be slightly better than the other choices.

**FIG. S2.**
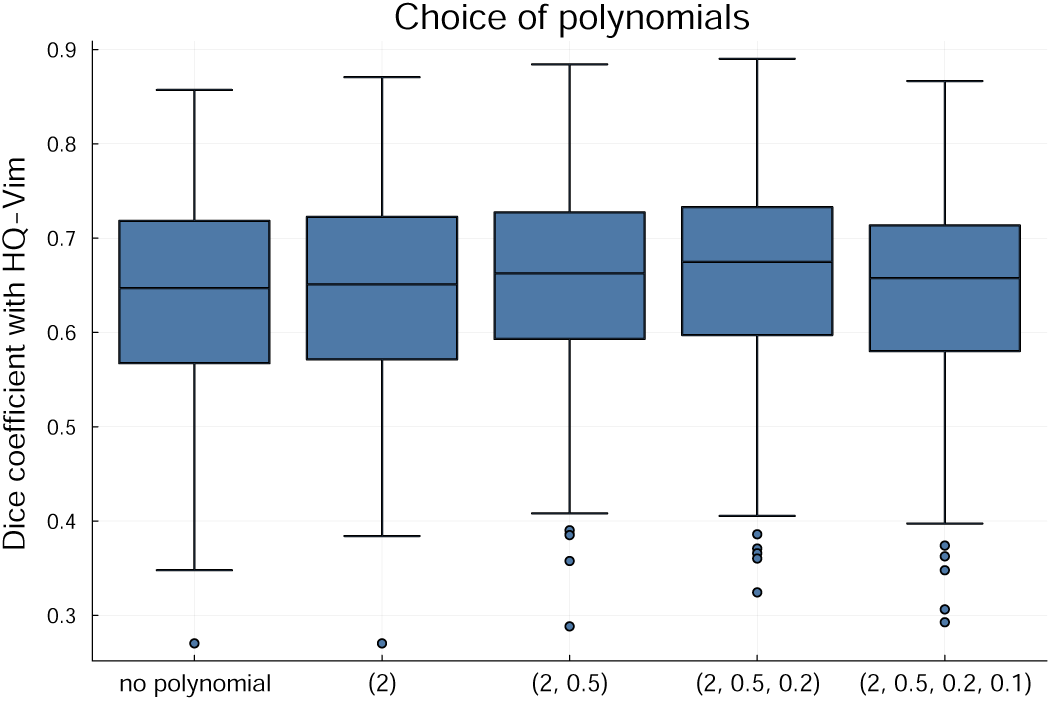
Choices of polynomial features. In addition to the original features (no polynomial), we tested a range of polynomial features on a smaller independent subset of subjects. The numbers in the x-axis denote the powers of polynomials used in the HQ-augmentation model, e.g., (2, 0.5, 0.2, 0.1) indicates that four additional polynomial features were included, with *p*1 = 2*, p*2 = 0.5*, p*3 = 0.2*, p*4 = 0.1 where *p*1*, p*2*, p*3*, p*4 were the powers. Overall, the choices of polynomials did not affect the Dice coefficient with HQ-Vim very much. However, (2, 0.5, 0.2), i.e., *p*1 = 2*, p*2 = 0.5*, p*3 = 0.2, appeared to be slightly better than the other choices.

**FIG. S3.**
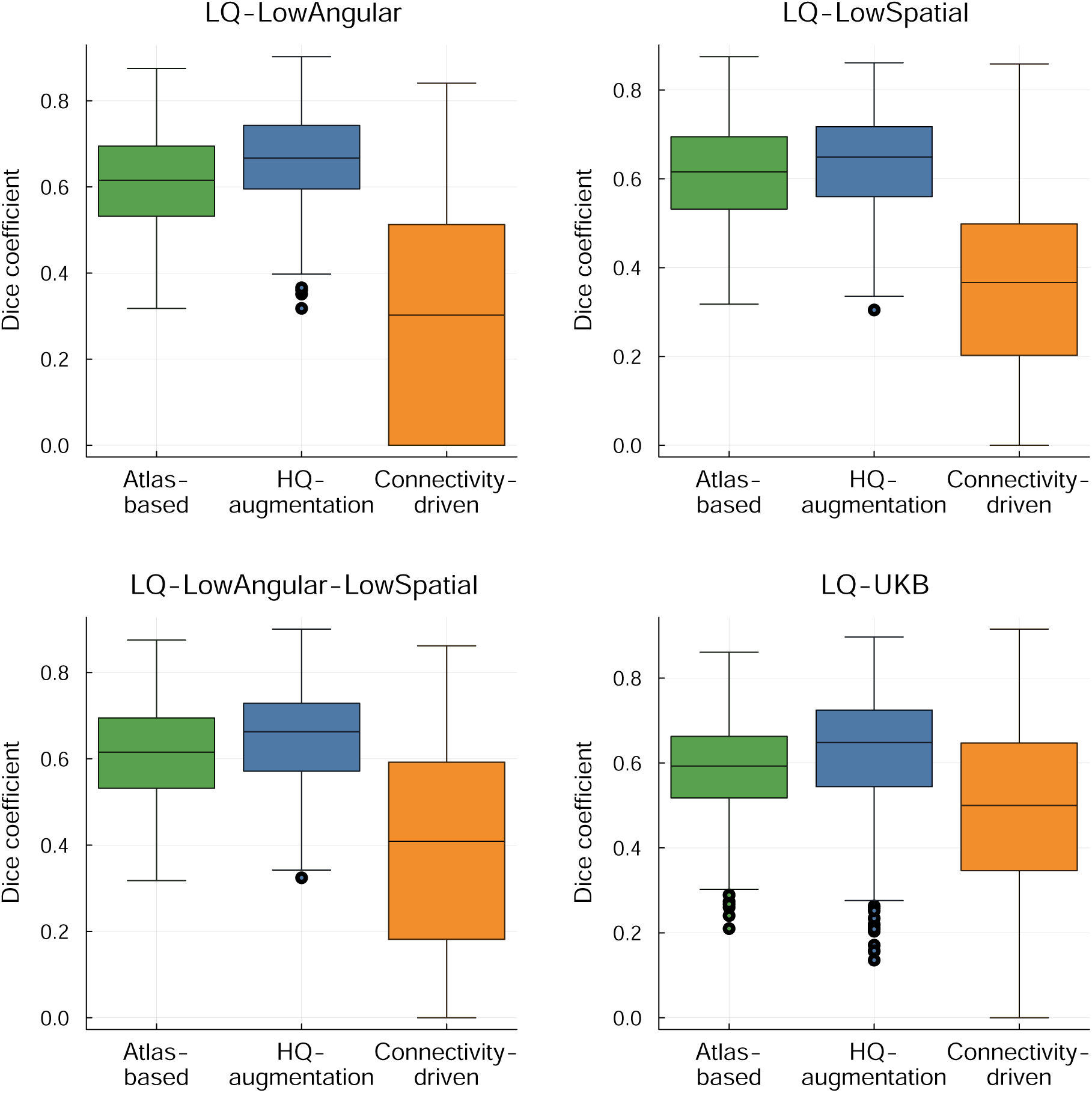
Boxplots of the Dice coefficient with HQ-Vim on surrogate low-quality datasets. **(A)**. Dice coefficient with the HQ-Vim, produced by the atlas-based (green), HQ-augmentation (blue), and connectivity-driven (orange) approach, on *LQ-LowAngular*. **(B)**. Equivalent plots of **(A)**, on *LQ-LowSpatial*. **(C)**. Equivalent plots of **(A)**, on *LQ-LowAngular-LowSpatial*. **(D)**. Equivalent plots of **(A)**, on LQ-UKB (where the HQ-augmentation model was trained on HCP and applied on LQ-UKB).

**FIG. S4.**
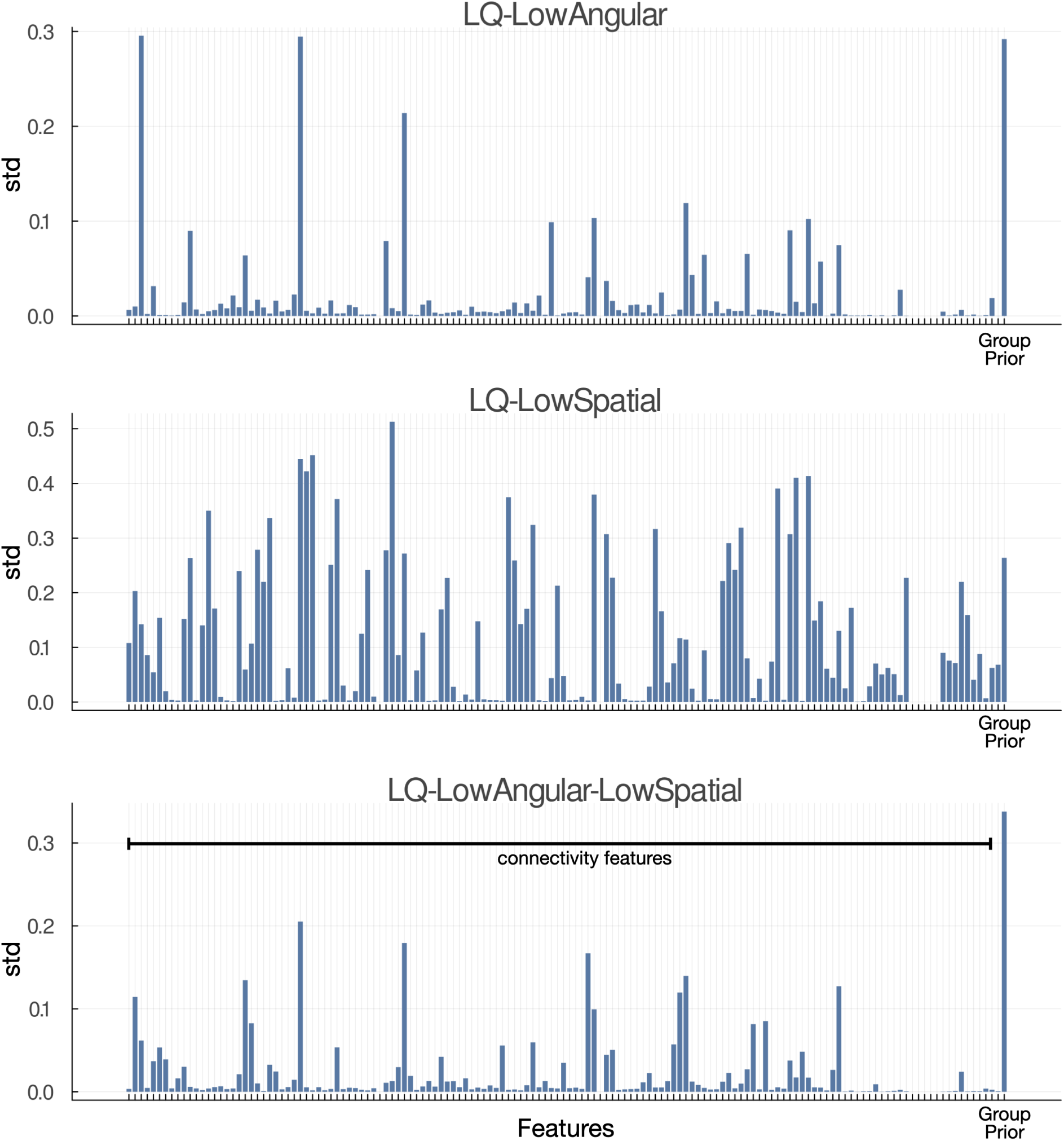
Relative contributions of the features in HQ-augmentation model for *LQ-LowAngular*, *LQ-LowSpatial*, *LQ-LowAngular-LowSpatial*. We plotted the std. of each feature contribution (i.e., the feature times its coefficient) across 50 subjects. The group-average prior feature contributed the most to Vim prediction, which is unsurprising given that the ground truth HQ-Vim labels were confined to be close to the atlas-defined Vim. For the purpose of visualisation, we summed the std. across the polynomials of a given connectivity feature. For example, the std. for M1 shown here was obtained by summing the std. of M1’s polynomial features. Specifically, suppose *β*1 is the coefficient for **x**M1, and *β*2 for **x**^2^, *β*3 for **x**^0.5^, and *β*4 for **x**^0.2^; then the std for M1 shown here is *std*(*β*1**x**M1) + *std*(*β*2**x**^2^) + *std*(*β*3**x**^0.5^) + *std*(*β*4**x**^0.2^) across 50 subjects.

### LISt of rois

We provide a table listing the ROIs used in this study, as well as the respective tractography protocols and the procedures to obtain them.

**TABLE S1:**
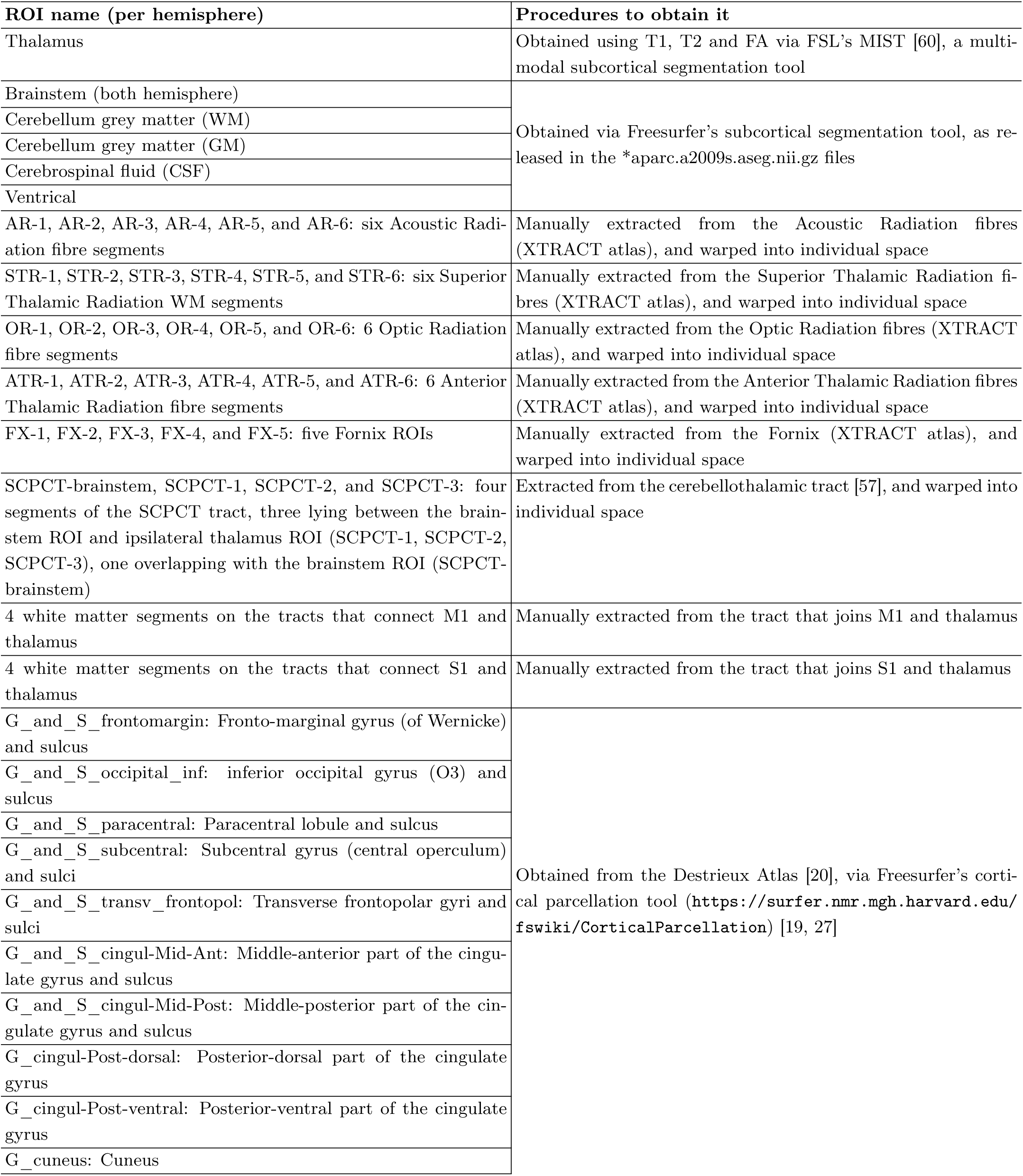
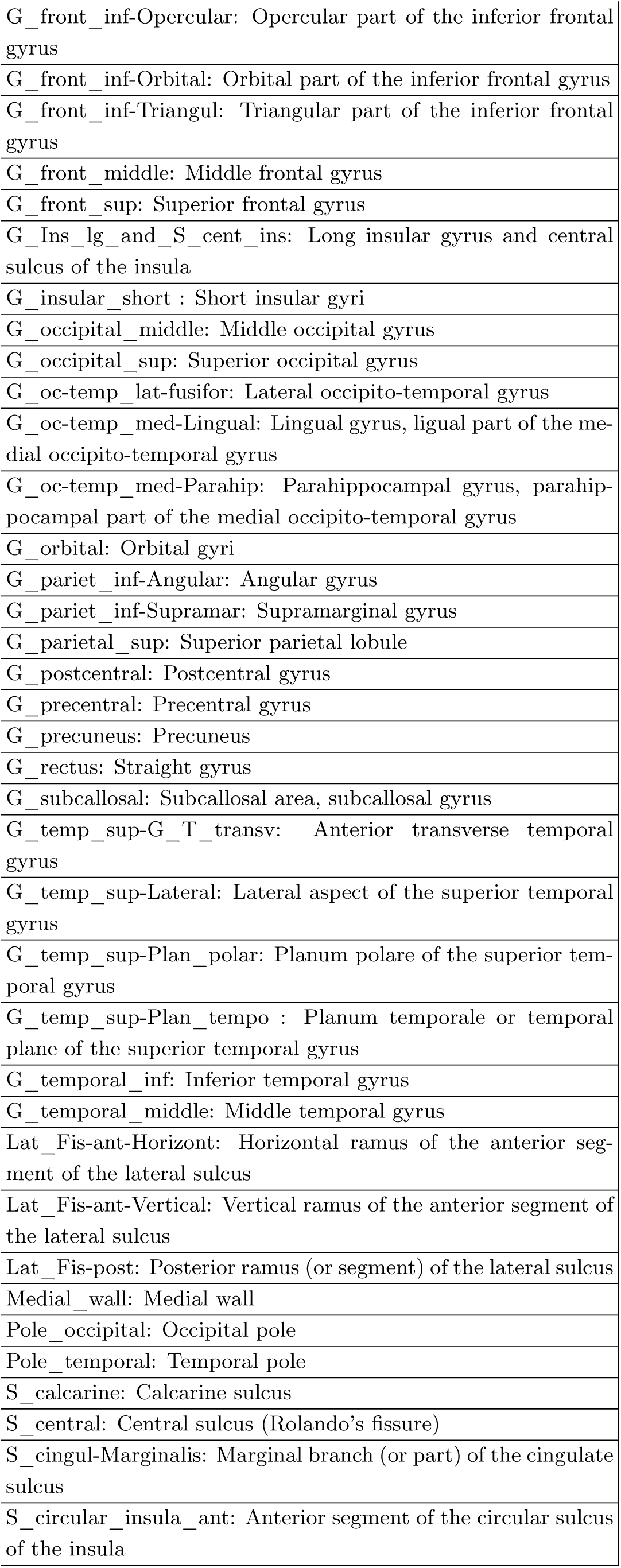
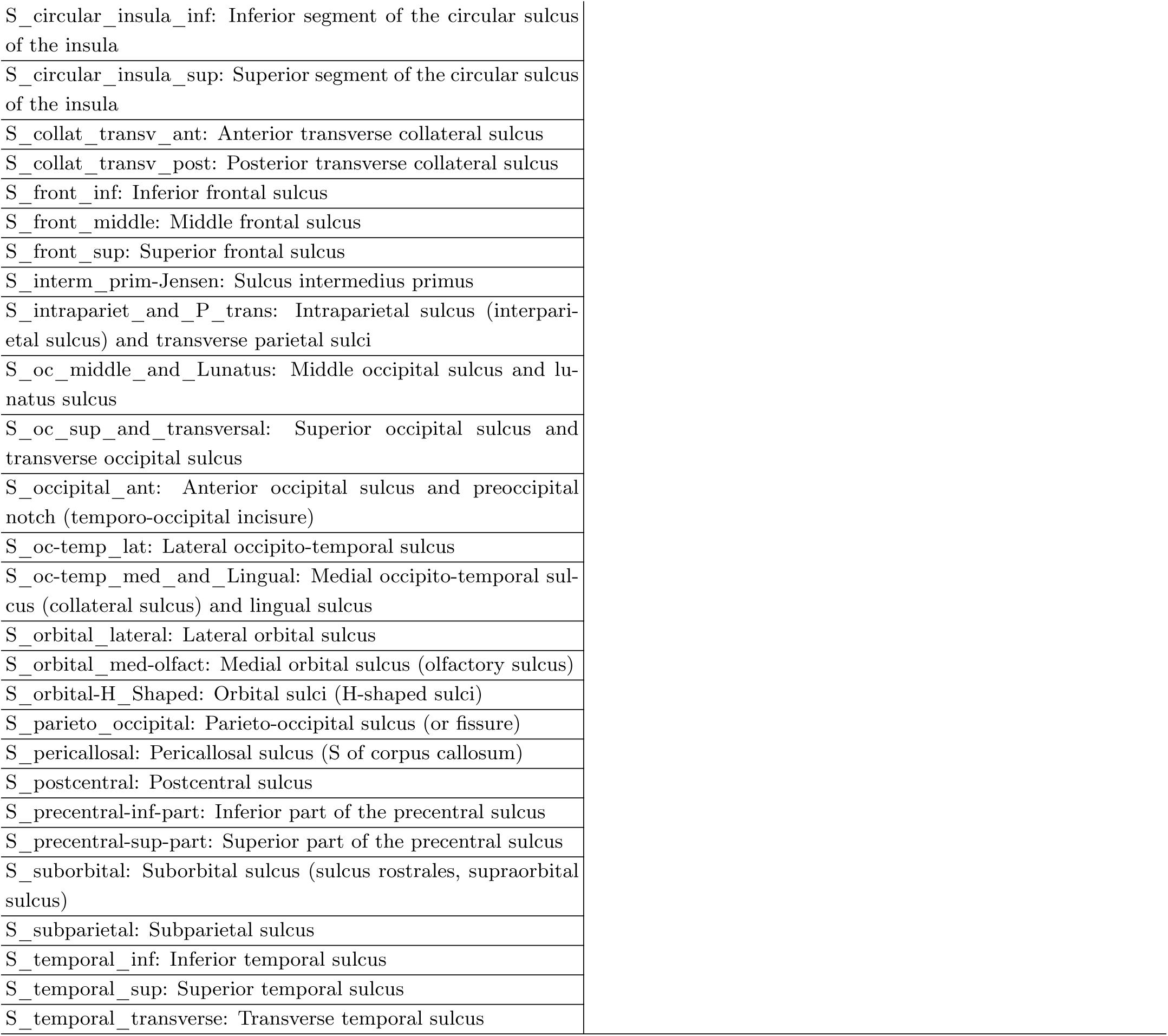
The list of anatomical ROIs used in this study.

| ROI name (per hemisphere) | Procedures to obtain it |
| --- | --- |
| Thalamus | Obtained using T1, T2 and FA via FSL's MIST [60], a multi-modal subcortical segmentation tool |
| Brainstem (both hemisphere) | Obtained via Freesurfer's subcortical segmentation tool, as released in the *aparc.a2009s.aseg.nii.gz files |
| Cerebellum grey matter (WM) |  |
| Cerebellum grey matter (GM) |  |
| Cerebrospinal fluid (CSF) |  |
| Ventricular |  |
| AR-1, AR-2, AR-3, AR-4, AR-5, and AR-6: six Acoustic Radiation fibre segments | Manually extracted from the Acoustic Radiation fibres (XTRACT atlas), and warped into individual space |
| STR-1, STR-2, STR-3, STR-4, STR-5, and STR-6: six Superior Thalamic Radiation WM segments | Manually extracted from the Superior Thalamic Radiation fibres (XTRACT atlas), and warped into individual space |
| OR-1, OR-2, OR-3, OR-4, OR-5, and OR-6: 6 Optic Radiation fibre segments | Manually extracted from the Optic Radiation fibres (XTRACT atlas), and warped into individual space |
| ATR-1, ATR-2, ATR-3, ATR-4, ATR-5, and ATR-6: 6 Anterior Thalamic Radiation fibre segments | Manually extracted from the Anterior Thalamic Radiation fibres (XTRACT atlas), and warped into individual space |
| FX-1, FX-2, FX-3, FX-4, and FX-5: five Fornix ROIs | Manually extracted from the Fornix (XTRACT atlas), and warped into individual space |
| SCPCT-brainstem, SCPCT-1, SCPCT-2, and SCPCT-3: four segments of the SCPCT tract, three lying between the brainstem ROI and ipsilateral thalamus ROI (SCPCT-1, SCPCT-2, SCPCT-3), one overlapping with the brainstem ROI (SCPCT-brainstem) | Extracted from the cerebellothalamic tract [57], and warped into individual space |
| 4 white matter segments on the tracts that connect M1 and thalamus | Manually extracted from the tract that joins M1 and thalamus |
| 4 white matter segments on the tracts that connect S1 and thalamus | Manually extracted from the tract that joins S1 and thalamus |
| G_and_S_frontomargin: Fronto-marginal gyrus (of Wernicke) and sulcus | Obtained from the Destrieux Atlas [20], via Freesurfer's cortical parcellation tool (https://surfer.nmr.mgh.harvard.edu/fswiki/CorticalParcellation) [19, 27] |
| G_and_S_occipital_inf: inferior occipital gyrus (O3) and sulcus |  |
| G_and_S_paracentral: Paracentral lobule and sulcus |  |
| G_and_S_subcentral: Subcentral gyrus (central operculum) and sulci |  |
| G_and_S_transv_frontopol: Transverse frontopolar gyri and sulci |  |
| G_and_S_cingul-Mid-Ant: Middle-anterior part of the cingulate gyrus and sulcus |  |
| G_and_S_cingul-Mid-Post: Middle-posterior part of the cingulate gyrus and sulcus |  |
| G_cingul-Post-dorsal: Posterior-dorsal part of the cingulate gyrus |  |
| G_cingul-Post-ventral: Posterior-ventral part of the cingulate gyrus |  |
| G_cuneus: Cuneus |  |

|  |
| --- |
| G_front_inf-Opercular: Opercular part of the inferior frontal gyrus |
| G_front_inf-Orbital: Orbital part of the inferior frontal gyrus |
| G_front_inf-Triangul: Triangular part of the inferior frontal gyrus |
| G_front_middle: Middle frontal gyrus |
| G_front_sup: Superior frontal gyrus |
| G_Ins_lg_and_S_cent_ins: Long insular gyrus and central sulcus of the insula |
| G_insular_short : Short insular gyri |
| G_occipital_middle: Middle occipital gyrus |
| G_occipital_sup: Superior occipital gyrus |
| G_oc-temp_lat-fusifor: Lateral occipito-temporal gyrus |
| G_oc-temp_med-Lingual: Lingual gyrus, ligual part of the medial occipito-temporal gyrus |
| G_oc-temp_med-Parahip: Parahippocampal gyrus, parahippocampal part of the medial occipito-temporal gyrus |
| G_orbital: Orbital gyri |
| G_pariet_inf-Angular: Angular gyrus |
| G_pariet_inf-Supramar: Supramarginal gyrus |
| G_parietal_sup: Superior parietal lobule |
| G_postcentral: Postcentral gyrus |
| G_precentral: Precentral gyrus |
| G_precuneus: Precuneus |
| G_rectus: Straight gyrus |
| G_subcallosal: Subcallosal area, subcallosal gyrus |
| G_temp_sup-G_T_transv: Anterior transverse temporal gyrus |
| G_temp_sup-Lateral: Lateral aspect of the superior temporal gyrus |
| G_temp_sup-Plan_polar: Planum polare of the superior temporal gyrus |
| G_temp_sup-Plan_tempo : Planum temporale or temporal plane of the superior temporal gyrus |
| G_temporal_inf: Inferior temporal gyrus |
| G_temporal_middle: Middle temporal gyrus |
| Lat_Fis-ant-Horizont: Horizontal ramus of the anterior segment of the lateral sulcus |
| Lat_Fis-ant-Vertical: Vertical ramus of the anterior segment of the lateral sulcus |
| Lat_Fis-post: Posterior ramus (or segment) of the lateral sulcus |
| Medial_wall: Medial wall |
| Pole_occipital: Occipital pole |
| Pole_temporal: Temporal pole |
| S_calcarine: Calcarine sulcus |
| S_central: Central sulcus (Rolando's fissure) |
| S_cingul-Marginalis: Marginal branch (or part) of the cingulate sulcus |
| S_circular_insula_ant: Anterior segment of the circular sulcus of the insula |
| S_circular_insula_inf: Inferior segment of the circular sulcus of the insula |
| S_circular_insula_sup: Superior segment of the circular sulcus of the insula |
| S_collat_transv_ant: Anterior transverse collateral sulcus |
| S_collat_transv_post: Posterior transverse collateral sulcus |
| S_front_inf: Inferior frontal sulcus |
| S_front_middle: Middle frontal sulcus |
| S_front_sup: Superior frontal sulcus |
| S_interm_prim-Jensen: Sulcus intermedius primus |
| S_intrapariet_and_P_trans: Intraparietal sulcus (interparietal sulcus) and transverse parietal sulci |
| S_oc_middle_and_Lunatus: Middle occipital sulcus and lunatus sulcus |
| S_oc_sup_and_transversal: Superior occipital sulcus and transverse occipital sulcus |
| S_occipital_ant: Anterior occipital sulcus and preoccipital notch (temporo-occipital incisure) |
| S_oc-temp_lat: Lateral occipito-temporal sulcus |
| S_oc-temp_med_and_Lingual: Medial occipito-temporal sulcus (collateral sulcus) and lingual sulcus |
| S_orbital_lateral: Lateral orbital sulcus |
| S_orbital_med-olfact: Medial orbital sulcus (olfactory sulcus) |
| S_orbital-H_Shaped: Orbital sulci (H-shaped sulci) |
| S_parieto_occipital: Parieto-occipital sulcus (or fissure) |
| S_pericallosal: Pericallosal sulcus (S of corpus callosum) |
| S_postcentral: Postcentral sulcus |
| S_precentral-inf-part: Inferior part of the precentral sulcus |
| S_precentral-sup-part: Superior part of the precentral sulcus |
| S_suborbital: Suborbital sulcus (sulcus rostrales, supraorbital sulcus) |
| S_subparietal: Subparietal sulcus |
| S_temporal_inf: Inferior temporal sulcus |
| S_temporal_sup: Superior temporal sulcus |
| S_temporal_transverse: Transverse temporal sulcus |

### TRACTOGRAPHY PROTOCOLS

**TABLE S2:**
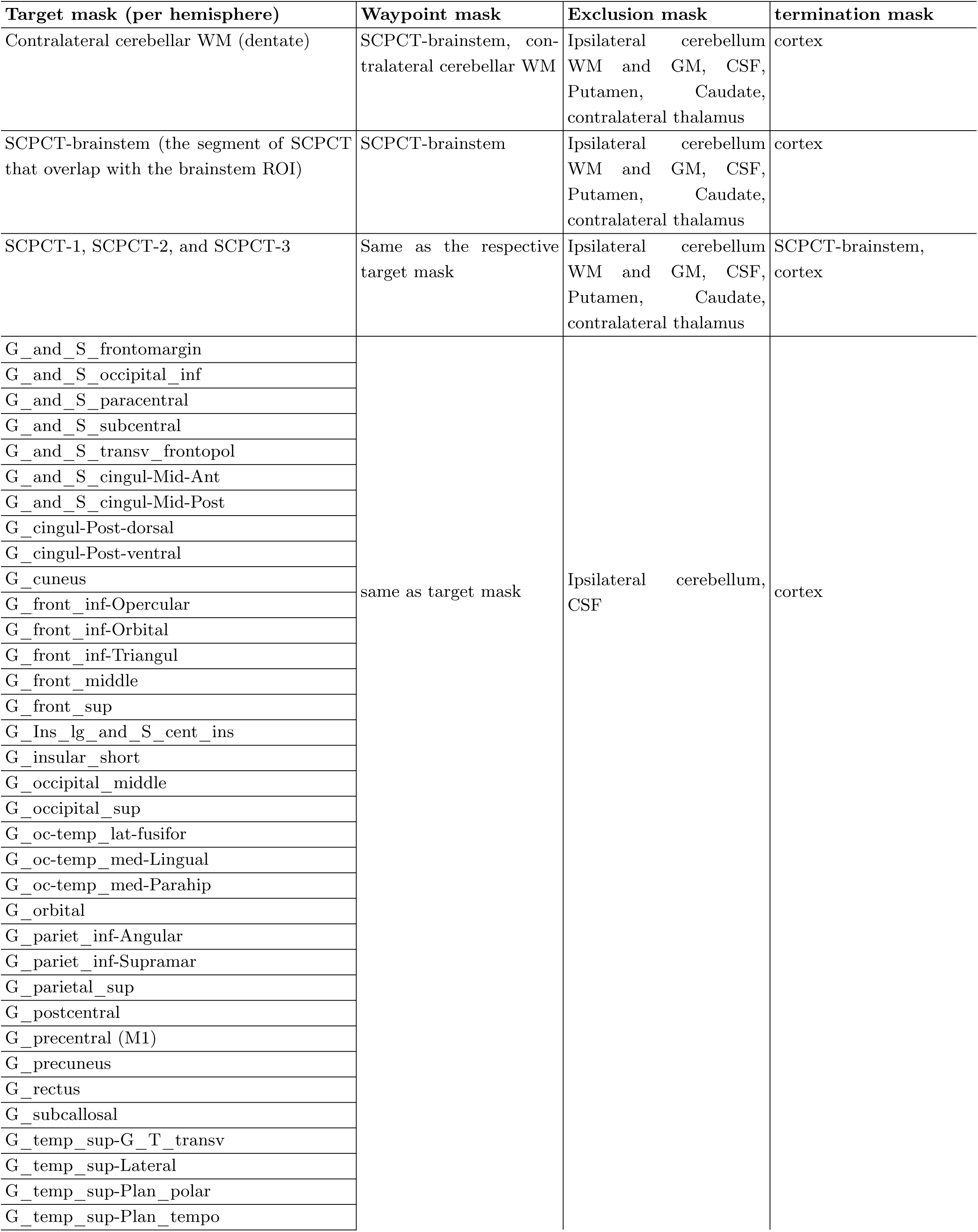
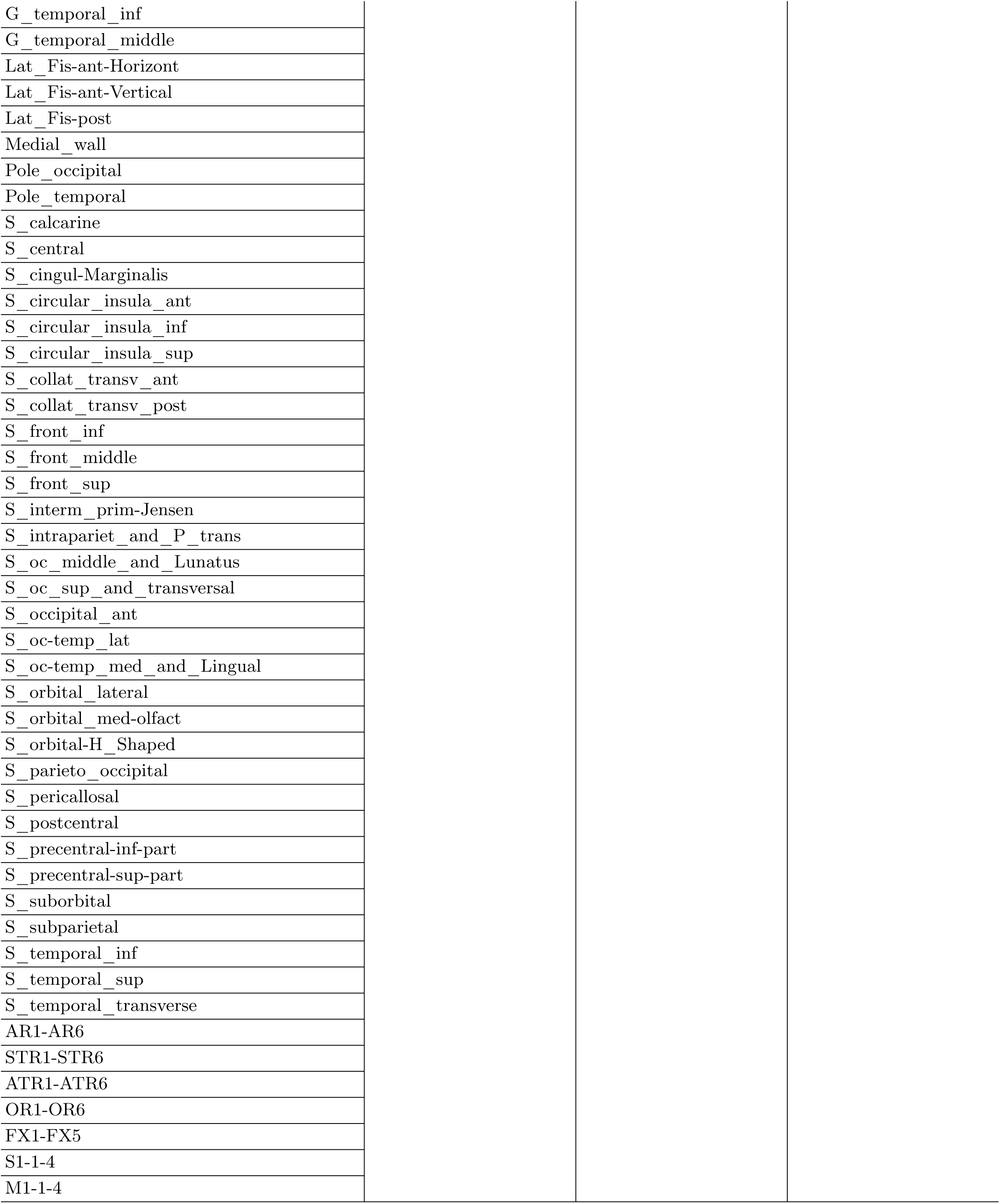
The list of seed, waypoint, exclusion and termination masks used in tractography.

| Target mask (per hemisphere) | Waypoint mask | Exclusion mask | termination mask |
| --- | --- | --- | --- |
| Contralateral cerebellar WM (dentate) | SCPCT-brainstem, contralateral cerebellar WM | Ipsilateral cerebellum WM and GM, CSF, Putamen, Caudate, contralateral thalamus | cortex |
| SCPCT-brainstem (the segment of SCPCT that overlap with the brainstem ROI) | SCPCT-brainstem | Ipsilateral cerebellum WM and GM, CSF, Putamen, Caudate, contralateral thalamus | cortex |
| SCPCT-1, SCPCT-2, and SCPCT-3 | Same as the respective target mask | Ipsilateral cerebellum WM and GM, CSF, Putamen, Caudate, contralateral thalamus | SCPCT-brainstem, cortex |
| G_and_S_frontomargin | same as target mask | Ipsilateral cerebellum, CSF | cortex |
| G_and_S_occipital_inf |  |  |  |
| G_and_S_paracentral |  |  |  |
| G_and_S_subcentral |  |  |  |
| G_and_S_transv_frontopol |  |  |  |
| G_and_S_cingul-Mid-Ant |  |  |  |
| G_and_S_cingul-Mid-Post |  |  |  |
| G_cingul-Post-dorsal |  |  |  |
| G_cingul-Post-ventral |  |  |  |
| G_cuneus |  |  |  |
| G_front_inf-Opercular |  |  |  |
| G_front_inf-Orbital |  |  |  |
| G_front_inf-Triangul |  |  |  |
| G_front_middle |  |  |  |
| G_front_sup |  |  |  |
| G_Ins_lg_and_S_cent_ins |  |  |  |
| G_insular_short |  |  |  |
| G_occipital_middle |  |  |  |
| G_occipital_sup |  |  |  |
| G_oc-temp_lat-fusifor |  |  |  |
| G_oc-temp_med-Lingual |  |  |  |
| G_oc-temp_med-Parahip |  |  |  |
| G_orbital |  |  |  |
| G_pariet_inf-Angular |  |  |  |
| G_pariet_inf-Supramar |  |  |  |
| G_parietal_sup |  |  |  |
| G_postcentral |  |  |  |
| G_precentral (M1) |  |  |  |
| G_precuneus |  |  |  |
| G_rectus |  |  |  |
| G_subcallosal |  |  |  |
| G_temp_sup-G_T_transv |  |  |  |
| G_temp_sup-Lateral |  |  |  |
| G_temp_sup-Plan_polar |  |  |  |
| G_temp_sup-Plan_tempo |  |  |  |

|  |
| --- |
| G_temporal_inf |
| G_temporal_middle |
| Lat_Fis-ant-Horizont |
| Lat_Fis-ant-Vertical |
| Lat_Fis-post |
| Medial_wall |
| Pole_occipital |
| Pole_temporal |
| S_calcarine |
| S_central |
| S_cingul-Marginalis |
| S_circular_insula_ant |
| S_circular_insula_inf |
| S_circular_insula_sup |
| S_collat_transv_ant |
| S_collat_transv_post |
| S_front_inf |
| S_front_middle |
| S_front_sup |
| S_interm_prim-Jensen |
| S_intrapariet_and_P_trans |
| S_oc_middle_and_Lunatus |
| S_oc_sup_and_transversal |
| S_occipital_ant |
| S_oc-temp_lat |
| S_oc-temp_med_and_Lingual |
| S_orbital_lateral |
| S_orbital_med-olfact |
| S_orbital-H_Shaped |
| S_parieto_occipital |
| S_pericallosal |
| S_postcentral |
| S_precentral-inf-part |
| S_precentral-sup-part |
| S_suborbital |
| S_subparietal |
| S_temporal_inf |
| S_temporal_sup |
| S_temporal_transverse |
| AR1-AR6 |
| STR1-STR6 |
| ATR1-ATR6 |
| OR1-OR6 |
| FX1-FX5 |
| S1-1-4 |
| M1-1-4 |

### MEAN-FIELD APPROXIMATION OF THE CRF DISTRIBUTION

As mentioned in section II F 2, we seek to maximise the probability of reproducing the exact same HQ-Vim label assignment **y** on its low-quality counterparts by minimising the cross entropy in Equation (1). Due to the inter-dependency of neighbouring voxels, the exact analytical minimisation is intractable. Thus, we approximate the CRF distribution *P* (**y***|***X**) by a simpler function *Q*(**y**), and iteratively solve this optimisation problem, as explained below. Suppose we have *K* classes (here *K* = 2). To initialise the model, we derived the initial coefficients 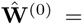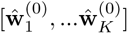 by optimising the following

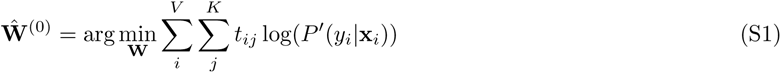

where *P^′^(y_i_|**x**_i_)* is the likelihood without considering local smoothness of the label assignment, i.e., ignoring the pairwise loss term in Equation (S1):

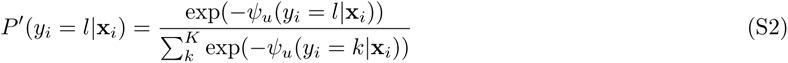

where, as defined before, 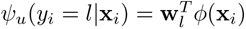. The coefficients 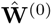 were used to initialise *Q*(**y**) using Equation (S2), i.e., *Q_i_(y_i_ = l) ← P^′^(y_i_ = l|**x**_i_)* evaluated at 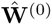 After initialisation, *Q_i_*(*y_i_*= *l*) is updated as the weighted sum of its neighbouring Q values, 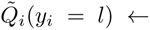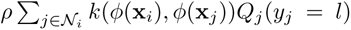, where *Q_i_*(*y_i_* = *l*) is the updated Q value. This is the message-passing step and is equivalent to applying M image-dependent Gaussian filters on the Q values. After message passing, label incompatibility was calculated as a penalty to encourage local smoothness. The incompatibility for label *l* at a given voxel *i*, denoted by *Q*^^^*_i_*(*y_i_* = *l*), was calculated as the sum of the updated *Q_i_* that takes a different label, i.e., 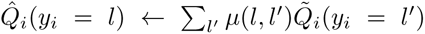. Next, this penalty incurred by incompatibility was subtracted from the unary inputs 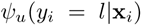, i.e., 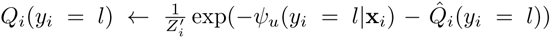 where 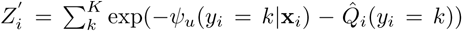 is the normalisation constant. The above steps were repeated until *Q* converges. The resulting *Q_i_*(*y_i_|***x***_i_*) is an approximation of the likelihood *P* (*y_i_|***x***_i_*), and was used when calculating the cross entropy in Equation (1). This cross entropy (1) was minimised in a mini-batch style via an ADAM optimiser [44] with learning rate 0.01, in which the connectivity features **X** of each subject served as a mini-batch. The model was trained using only the reliable subjects. To clarify, “reliable” refers to the subjects that passed the four selection criteria. These reliable subjects were further divided into two separate groups or “folds”. The model was trained using one fold at a time, with the other fold and “unreliable” subjects (those who did not pass the quality control) used for testing. This approach ensured that our model was exposed to and learned from the most accurate data available.

The pseudo code of the above steps is summarised in Table S3.

**TABLE S3.**
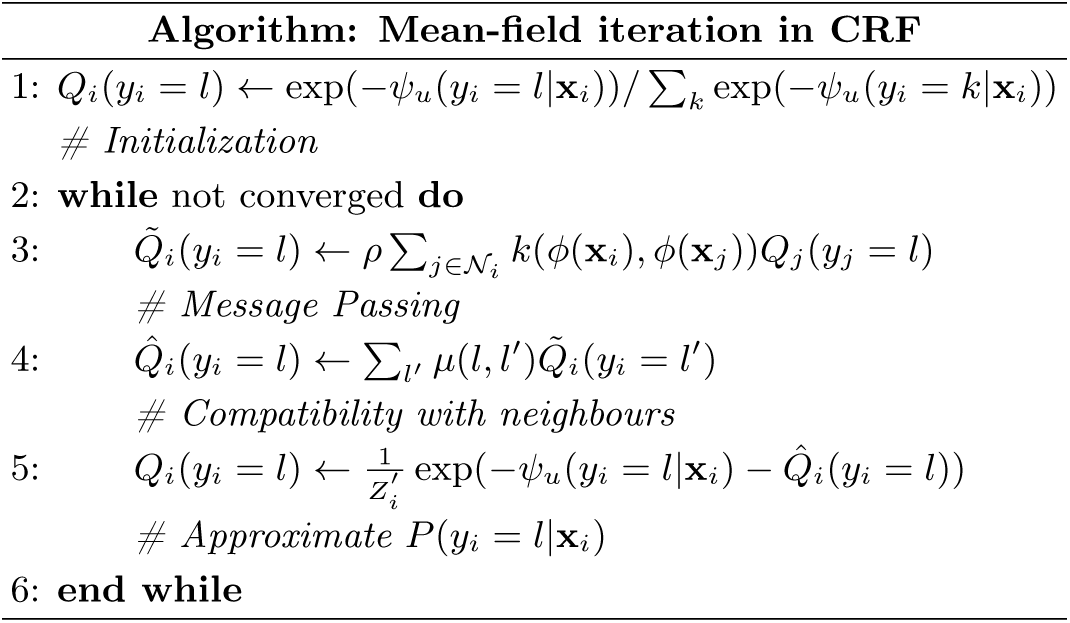
Pseudo code of the mean-field iteration to solve *Q*(**y**), an approximation of the CRF distribution *P* (**y**|**X**).

**TABLE S4.** P values of paired t-tests comparing the HQ-augmentation against the alternative methods, based on their correspondence with HQ-Vim. Metrics used for comparison include the Dice coefficient and centroid displacement. P values were Bonferroni corrected (16 tests in total).

|  | *LQ-LowAngular* |  | *LQ-LowSpatial* |  |
| --- | --- | --- | --- | --- |
|  | Dice<br>coefficient | Centroid<br>displacement | Dice<br>coefficient | Centroid<br>displacement |
| HQ-augmentation<br>vs connectivity-driven | <1e-99 | <1e-54 | <1e-99 | <1e-89 |
| HQ-augmentation<br>vs atlas-based | <1e-10 | <1e-17 | 0.0043 | <1e-17 |
|  | *LQ-LowAngular-LowSpatial* |  | LQ-UKB |  |
|  | Dice<br>coefficient | Centroid<br>displacement | Dice<br>coefficient | Centroid<br>displacement |
| HQ-augmentation<br>vs connectivity-driven | <1e-72 | <1e-38 | <1e-96 | <1e-66 |
| HQ-augmentation<br>vs atlas-based | <1e-4 | 1e-12 | <1e-19 | <1e-12 |

## References

[1] Akram, H., Dayal, V., Mahlknecht, P., Georgiev, D., Hyam, J., Foltynie, T., Limousin, P., De Vita, E., Jahanshahi, M., Ashburner, J., et al. (2018). Connectivity derived thalamic segmentation in deep brain stimulation for tremor. NeuroImage: Clinical, 18:130–142.

[2] Alexander, D. C., Zikic, D., Ghosh, A., Tanno, R., Wottschel, V., Zhang, J., Kaden, E., Dyrby, T. B., Sotiropoulos, S. N., Zhang, H., et al. (2017). Image quality transfer and applications in diffusion mri. NeuroImage, 152:283–298.

[3] Alexander, D. C., Zikic, D., Zhang, J., Zhang, H., and Criminisi, A. (2014). Image quality transfer via random forest regression: applications in diffusion mri. In Medical Image Computing and Computer-Assisted Intervention– MICCAI 2014: 17th International Conference, Boston, MA, USA, September 14-18, 2014, Proceedings, Part III 17, pages 225–232. Springer.

[4] Alfaro-Almagro, F., Jenkinson, M., Bangerter, N. K., Andersson, J. L., Griffanti, L., Douaud, G., Sotiropoulos, S. N., Jbabdi, S., Hernandez-Fernandez, M., Vallee, E., et al. (2018). Image processing and quality control for the first 10,000 brain imaging datasets from uk biobank. Neuroimage, 166:400–424.

[5] Anderson, V. C., Burchiel, K. J., Hogarth, P., Favre, J., and Hammerstad, J. P. (2005). Pallidal vs subthalamic nucleus deep brain stimulation in parkinson disease. Archives of Neurology, 62(4):554–560.

[6] Andersson, J. L., Jenkinson, M., and Smith, S. (2007). Non-linear optimisation fmrib technical report tr07ja1. University of Oxford FMRIB Centre: Oxford UK.

[7] Andersson, J. L., Skare, S., and Ashburner, J. (2003). How to correct susceptibility distortions in spin-echo echo-planar images: application to diffusion tensor imaging. Neuroimage, 20(2):870–888.

[8] Andersson, J. L. and Sotiropoulos, S. N. (2016). An integrated approach to correction for off-resonance effects and subject movement in diffusion mr imaging. Neuroimage, 125:1063–1078.

[9] Bailey, P. (1959). Introduction to Stereotaxis with an Atlas of Human Brain: In Three Volumes. Georg Thieme Verlag.

[10] Baker, K. B., Schuster, D., Cooperrider, J., and Machado, A. G. (2010). Deep brain stimulation of the lateral cerebellar nucleus produces frequency-specific alterations in motor evoked potentials in the rat in vivo. Experimental neurology, 226(2):259–264.

[11] Basser, P. J., Mattiello, J., and LeBihan, D. (1994). Estimation of the effective self-diffusion tensor from the nmr spin echo. Journal of Magnetic Resonance, Series B, 103(3):247–254.

[12] Behrens, T. E., Berg, H. J., Jbabdi, S., Rushworth, M. F., and Woolrich, M. W. (2007). Probabilistic diffusion tractography with multiple fibre orientations: What can we gain? neuroimage, 34(1):144–155.

[13] Behrens, T. E., Woolrich, M. W., Jenkinson, M., Johansen-Berg, H., Nunes, R. G., Clare, S., Matthews, P. M., Brady, J. M., and Smith, S. M. (2003). Characterization and propagation of uncertainty in diffusion-weighted mr imaging. Magnetic Resonance in Medicine: An Official Journal of the International Society for Magnetic Resonance in Medicine, 50(5):1077–1088.

[14] Benabid, A.-L., Chabardes, S., Mitrofanis, J., and Pollak, P. (2009). Deep brain stimulation of the subthalamic nucleus for the treatment of parkinson’s disease. The Lancet Neurology, 8(1):67–81.

[15] Bertino, S., Basile, G. A., Bramanti, A., Ciurleo, R., Tisano, A., Anastasi, G. P., Milardi, D., and Cacciola, A. (2021). Ventral intermediate nucleus structural connectivity-derived segmentation: anatomical reliability and variability. Neuroimage, 243:118519.

[16] Blumberg, S. B., Tanno, R., Kokkinos, I., and Alexander, D. C. (2018). Deeper image quality transfer: Training low-memory neural networks for 3d images. In Medical Image Computing and Computer Assisted Intervention– MICCAI 2018: 21st International Conference, Granada, Spain, September 16-20, 2018, Proceedings, Part I, pages 118–125. Springer.

[17] Calzavara, R., Zappalà, A., Rozzi, S., Matelli, M., and Luppino, G. (2005). Neurochemical characterization of the cerebellar-recipient motor thalamic territory in the macaque monkey. European Journal of Neuroscience, 21(7):1869–1894.

[18] Cury, R. G., Fraix, V., Castrioto, A., Fernández, M. A. P., Krack, P., Chabardes, S., Seigneuret, E., Alho, E. J. L., Benabid, A.-L., and Moro, E. (2017). Thalamic deep brain stimulation for tremor in parkinson disease, essential tremor, and dystonia. Neurology, 89(13):1416– 1423.

[19] Desikan, R. S., Ségonne, F., Fischl, B., Quinn, B. T., Dickerson, B. C., Blacker, D., Buckner, R. L., Dale, A. M., Maguire, R. P., Hyman, B. T., et al. (2006). An automated labeling system for subdividing the human cerebral cortex on mri scans into gyral based regions of interest. Neuroimage, 31(3):968–980.

[20] Destrieux, C., Fischl, B., Dale, A., and Halgren, E. (2010). Automatic parcellation of human cortical gyri and sulci using standard anatomical nomenclature. Neuroimage, 53(1):1–15.

[21] Dum, R. P. and Strick, P. L. (2003). An unfolded map of the cerebellar dentate nucleus and its projections to the cerebral cortex. Journal of neurophysiology, 89(1):634– 639.

[22] Elias, G. J., Boutet, A., Joel, S. E., Germann, J., Gwun, D., Neudorfer, C., Gramer, R. M., Algarni, M., Paramanandam, V., Prasad, S., et al. (2021). Probabilistic mapping of deep brain stimulation: insights from 15 years of therapy. Annals of Neurology, 89(3):426–443.

[23] Elias, W. J., Huss, D., Voss, T., Loomba, J., Khaled, M., Zadicario, E., Frysinger, R. C., Sperling, S. A., Wylie, S., Monteith, S. J., et al. (2013). A pilot study of focused ultrasound thalamotomy for essential tremor. New England Journal of Medicine, 369(7):640–648.

[24] Ferreira, F., Akram, H., Ashburner, J., Zrinzo, L., Zhang, H., and Lambert, C. (2021). Ventralis intermedius nucleus anatomical variability assessment by mri structural connectivity. NeuroImage, 238:118231.

[25] Figini, M., Lin, H., Ogbole, G., Arco, F. D., Blumberg, S. B., Carmichael, D. W., Tanno, R., Kaden, E., Brown, B. J., Lagunju, I., et al. (2020). Image quality transfer enhances contrast and resolution of low-field brain mri in african paediatric epilepsy patients. arXiv preprint arXiv:2003.07216.

[26] Fischl, B., Salat, D. H., Busa, E., Albert, M., Dieterich, M., Haselgrove, C., Van Der Kouwe, A., Killiany, R., Kennedy, D., Klaveness, S., et al. (2002). Whole brain segmentation: automated labeling of neuroanatomical structures in the human brain. Neuron, 33(3):341–355.

[27] Fischl, B., Van Der Kouwe, A., Destrieux, C., Halgren, E., Ségonne, F., Salat, D. H., Busa, E., Seidman, L. J., Goldstein, J., Kennedy, D., et al. (2004). Automatically parcellating the human cerebral cortex. Cerebral cortex, 14(1):11–22.

[28] Fisher, R., Salanova, V., Witt, T., Worth, R., Henry, T., Gross, R., Oommen, K., Osorio, I., Nazzaro, J., Labar, D., et al. (2010). Electrical stimulation of the anterior nucleus of thalamus for treatment of refractory epilepsy. Epilepsia, 51(5):899–908.

[29] Gallay, M. N., Jeanmonod, D., Liu, J., and Morel, A. (2008). Human pallidothalamic and cerebellothalamic tracts: anatomical basis for functional stereotactic neurosurgery. Brain Structure and Function, 212:443–463.

[30] Glasser, M. F., Sotiropoulos, S. N., Wilson, J. A., Coalson, T. S., Fischl, B., Andersson, J. L., Xu, J., Jbabdi, S., Webster, M., Polimeni, J. R., et al. (2013). The minimal preprocessing pipelines for the human connectome project. Neuroimage, 80:105–124.

[31] Grabner, G., Janke, A. L., Budge, M. M., Smith, D., Pruessner, J., and Collins, D. L. (2006). Symmetric atlasing and model based segmentation: an application to the hippocampus in older adults. In International Conference on Medical Image Computing and Computer-Assisted Intervention, pages 58–66. Springer.

[32] Gravbrot, N., Saranathan, M., Pouratian, N., and Kasoff, W. S. (2020). Advanced imaging and direct targeting of the motor thalamus and dentato-rubro-thalamic tract for tremor: a systematic review. Stereotactic and Functional Neurosurgery, 98(4):220–240.

[33] Haq, I. U., Foote, K. D., Goodman, W. K., Ricciuti, N., Ward, H., Sudhyadhom, A., Jacobson, C. E., Siddiqui, M. S., and Okun, M. S. (2010). A case of mania following deep brain stimulation for obsessive compulsive disorder. Stereotactic and functional neurosurgery, 88(5):322–328.

[34] Helmich, R. C., Hallett, M., Deuschl, G., Toni, I., and Bloem, B. R. (2012). Cerebral causes and consequences of parkinsonian resting tremor: a tale of two circuits? Brain, 135(11):3206–3226.

[35] Hernandez-Fernandez, M., Reguly, I., Jbabdi, S., Giles, M., Smith, S., and Sotiropoulos, S. N. (2019). Using gpus to accelerate computational diffusion mri: From microstructure estimation to tractography and connectomes. Neuroimage, 188:598–615.

[36] Hirai, T. and Jones, E. (1989). A new parcellation of the human thalamus on the basis of histochemical staining. Brain Research Reviews, 14(1):1–34.

[37] Hodaie, M., Wennberg, R. A., Dostrovsky, J. O., and Lozano, A. M. (2002). Chronic anterior thalamus stimulation for intractable epilepsy. Epilepsia, 43(6):603–608.

[38] Jakab, A., Blanc, R., Berényi, E., and Székely, G. (2012). Generation of individualized thalamus target maps by using statistical shape models and thalamocortical tractography. American Journal of Neuroradiology, 33(11):2110–2116.

[39] Jbabdi, S., Sotiropoulos, S. N., Savio, A. M., Graña, M., and Behrens, T. E. (2012). Model-based analysis of multishell diffusion mr data for tractography: How to get over fitting problems. Magnetic resonance in medicine, 68(6):1846–1855.

[40] Jenkinson, M., Bannister, P., Brady, M., and Smith, S. (2002). Improved optimization for the robust and accurate linear registration and motion correction of brain images. Neuroimage, 17(2):825–841.

[41] Jenkinson, M., Beckmann, C. F., Behrens, T. E., Woolrich, M. W., and Smith, S. M. (2012). Fsl. Neuroimage, 62(2):782–790.

[42] Johansen-Berg, H., Behrens, T. E., Sillery, E., Ciccarelli, O., Thompson, A. J., Smith, S. M., and Matthews, P. M. (2005). Functional–anatomical validation and individual variation of diffusion tractography-based segmentation of the human thalamus. Cerebral cortex, 15(1):31–39.

[43] Jones, E. G. (2007). The Thalamus. Cambridge University Press, Cambridge.

[44] Kingma, D. P. and Ba, J. (2014). Adam: A method for stochastic optimization. arXiv preprint arXiv:1412.6980.

[45] Krack, P., Batir, A., Van Blercom, N., Chabardes, S., Fraix, V., Ardouin, C., Koudsie, A., Limousin, P. D., Benazzouz, A., LeBas, J.-F., Benabid, A.-L., and Pollak, P. (2003). Five-year follow-up of bilateral stimulation of the subthalamic nucleus in advanced parkinson’s disease. New England Journal of Medicine, 349(20):1925–1934.

[46] Lee, K. J., Shon, Y. M., and Cho, C. B. (2012). Long-term outcome of anterior thalamic nucleus stimulation for intractable epilepsy. Stereotactic and functional neurosurgery, 90(6):379–385.

[47] McIntyre, C. C. and Hahn, P. J. (2010). Network perspectives on the mechanisms of deep brain stimulation. Neurobiology of disease, 38(3):329–337.

[48] Miller, K. L., Alfaro-Almagro, F., Bangerter, N. K., Thomas, D. L., Yacoub, E., Xu, J., Bartsch, A. J., Jbabdi, S., Sotiropoulos, S. N., Andersson, J. L., et al. (2016). Multimodal population brain imaging in the uk biobank prospective epidemiological study. Nature neuroscience, 19(11):1523–1536.

[49] Morel, A., Magnin, M., and Jeanmonod, D. (1997). Multiarchitectonic and stereotactic atlas of the human thalamus. Journal of Comparative Neurology, 387(4):588–630.

[50] Odekerken, V. J., van Laar, T., Staal, M. J., Mosch, A., Hoffmann, C. F., Nijssen, P. C., Beute, G. N., van Vugt, J. P., Lenders, M. W., Contarino, M. F., et al. (2013). Subthalamic nucleus versus globus pallidus bilateral deep brain stimulation for advanced parkinson’s disease (nstaps study): a randomised controlled trial. The Lancet Neurology, 12(1):37–44.

[51] Peltola, J., Colon, A. J., Pimentel, J., Coenen, V. A., Gil-Nagel, A., Ferreira, A. G., Lehtimäki, K., Ryvlin, P., Taylor, R. S., Ackermans, L., et al. (2023). Deep brain stimulation of the anterior nucleus of the thalamus in drug resistant epilepsy in the more multicenter patient registry. Neurology.

[52] Rodriguez-Oroz, M. C., Obeso, J. A., Lang, A. E., Houeto, J.-L., Pollak, P., Rehncrona, S., Kulisevsky, J., Albanese, A., Volkmann, J., Hariz, M. I., et al. (2005). Bilateral deep brain stimulation in parkinson’s disease: a multicentre study with 4 years follow-up. Brain, 128(10):2240–2249.

[53] Sakai, S. T. (2021). Cerebellar thalamic and thalamocortical projections. In Handbook of the Cerebellum and Cerebellar Disorders, pages 661–680. Springer.

[54] Sammartino, F., Krishna, V., King, N. K. K., Lozano, A. M., Schwartz, M. L., Huang, Y., and Hodaie, M. (2016). Tractography-based ventral intermediate nucleus targeting: Novel methodology and intraoperative validation. Movement Disorders, 31(8):1217–1225.

[55] Sotiropoulos, S. N., Hernández-Fernández, M., Vu, A. T., Andersson, J. L., Moeller, S., Yacoub, E., Lenglet, C., Ugurbil, K., Behrens, T. E., and Jbabdi, S. (2016). Fusion in diffusion mri for improved fibre orientation estimation: An application to the 3t and 7t data of the human connectome project. Neuroimage, 134:396–409.

[56] Sotiropoulos, S. N., Jbabdi, S., Xu, J., Andersson, J. L., Moeller, S., Auerbach, E. J., Glasser, M. F., Hernandez, M., Sapiro, G., Jenkinson, M., et al. (2013). Advances in diffusion mri acquisition and processing in the human connectome project. Neuroimage, 80:125–143.

[57] Tang, Y., Sun, W., Toga, A. W., Ringman, J. M., and Shi, Y. (2018). A probabilistic atlas of human brainstem pathways based on connectome imaging data. Neuroimage, 169:227–239.

[58] Tanno, R., Worrall, D. E., Ghosh, A., Kaden, E., Sotiropoulos, S. N., Criminisi, A., and Alexander, D. C. (2017). Bayesian image quality transfer with cnns: exploring uncertainty in dmri super-resolution. In Medical Image Computing and Computer Assisted Intervention-MICCAI 2017: 20th International Conference, Quebec City, QC, Canada, September 11-13, 2017, Proceedings, Part I 20, pages 611–619. Springer.

[59] Van Essen, D. C., Smith, S. M., Barch, D. M., Behrens, T. E., Yacoub, E., Ugurbil, K., Consortium, W.-M. H., et al. (2013). The wu-minn human connectome project: an overview. Neuroimage, 80:62–79.

[60] Visser, E., Keuken, M. C., Douaud, G., Gaura, V., Bachoud-Levi, A.-C., Remy, P., Forstmann, B. U., and Jenkinson, M. (2016). Automatic segmentation of the striatum and globus pallidus using mist: Multimodal image segmentation tool. NeuroImage, 125:479–497.

[61] Volkmann, J., Allert, N., Voges, J., Weiss, P. H., Freund, H.-J., and Sturm, V. (2004). Safety and efficacy of pallidal or subthalamic nucleus stimulation in advanced pd. Neurology, 63(5):915–920.

[62] Warrington, S., Bryant, K. L., Khrapitchev, A. A., Sallet, J., Charquero-Ballester, M., Douaud, G., Jbabdi, S., Mars, R. B., and Sotiropoulos, S. N. (2020). Xtract-standardised protocols for automated tractography in the human and macaque brain. Neuroimage, 217:116923.

[63] Zheng, S., Jayasumana, S., Romera-Paredes, B., Vineet, V., Su, Z., Du, D., Huang, C., and Torr, P. H. (2015). Conditional random fields as recurrent neural networks. In Proceedings of the IEEE international conference on computer vision, pages 1529–1537.

[64] Zrinzo, L., Foltynie, T., Limousin, P., and Hariz, M. I. (2012). Reducing hemorrhagic complications in functional neurosurgery: a large case series and systematic literature review. Journal of neurosurgery, 116(1):84–94.

